# EpiMII: Integrating Structure and Graph Neural Networks for MHC-II Epitope and Neoantigen Design

**DOI:** 10.1101/2025.04.29.651031

**Authors:** Jiayi Yuan, Xiaowei Xu, Ze-Yu Sun, Tianjian Liang, Jingxuan Ge, Ruofan Jin, Xiang-Qun Xie, Yan Chen, Tingjun Hou, Zhiwei Feng

## Abstract

MHC-II neoantigens play a critical role in immunotherapy, either as direct effectors or through their influence on CD8^+^ T cells. However, only a small fraction of tumor DNA mutations qualify as functional neoantigens, and current prediction tools often lack accuracy, leading to the low immunogenicity of predicted neoantigens in vivo. Here, we present EpiMII, a Graph Neural Network model for MHC-II epitope design, which learns from the structural features of epitopes to predict their sequences. To train EpiMII, we constructed a reliable, large dataset containing 142,934 MHC-II epitope structures. This approach achieves a 4.2x improvement over ProteinMPNN, with a sequence recovery rate of 78.0% for known MHC-II epitopes in the Protein Data Bank. As a case study, we designed a neoantigen from hepatocellular carcinoma. All five designed epitopes significantly activated CD4^+^ T cells in vitro and induced secretion of IFN-γ and TNF-α. Notably, one epitope treatment significantly reduced tumor volume in mice in vivo. EpiMII offers a novel and efficient approach for identifying MHC-II epitopes/neoantigens, potentially contributing to vaccine development.

## Introduction

Neoantigens, derived from non-synonymous somatic mutations in tumor cells and absent in normal cells, are recognized as non-self antigens, evoking an immune response not subject to central and peripheral tolerance[1–3]. In the tumor microenvironment and periphery of human patients with cancer, neoantigen-specific T cells are frequently observed before and during treatment with immunotherapies such as immune checkpoint blockade and personalized cancer vaccines[4]. Early clinical trials of neoantigen-based therapeutic vaccines for standard cancer treatment have yielded encouraging outcomes in monotherapy or combination therapy with immune checkpoint inhibitors (ICIs)[5–7].

The critical roles of MHC-II-restricted CD4^+^ T cell responses to tumor neoantigens during immunotherapy have been revealed recently[8], either as direct effectors or through indirect influence on CD8^+^ T cells[4]. As the key to the activation of CD4^+^ T cells, MHC-II expression was observed in multiple human tumor cells, including melanoma, ovarian cancer, lung cancer, and more[9]. Besides, in the absence of tumor cell-intrinsic MHC-II expression, CD4^+^ T cells are still proved to help with killing tumors[10]. This outperforms the MHC-I antigen presentation pathway where MHC-I suffers the immune escape mechanism, significantly decreasing the efficacy of MHC-I epitopes. Most reports also demonstrate that CD4^+^ T cell responses to MHC-II epitope are not only required for optimal priming of MHC-I-restricted CD8^+^ T cells and their mutation into cytotoxic T cells, but also required for robust responses to ICIs[8, 11, 12]. Regulatory T cells (Treg) have been known to impede anti-tumor response. CD4^+^ T cells promote tumor rejection by inhibiting the response triggered by Treg when the vaccine contains both MHC-I neoantigens and low-dose MHC-II-restricted neoantigens[13]. All these findings make the MHC-II neoantigen the most promising target for tumor treatment. However, only a minor fraction of DNA mutations presents in cancer cells qualify as neoantigen. Among the identified mutations, most cancer mutations are unique to individual patients, and only a few are shared among some patients[1], which restricts the development of MHC-II-neoantigen-based cancer immunotherapy at the personal level but not at the population level.

Emerging tools that can be applied to MHC-II neoantigen identification of different cancer types mainly focus on 1) MHC-II binding affinity prediction, such as the widely used NetMHCIIpan4.0[14], and MixMHC2pred[15]; 2) Immunogenicity prediction, such as FIONA[16], TLimmuno2[17], DeepNeo-v2[18], and CD4episcore[19]. MARIA predicted the MHC-II epitope in both binding affinity and immunogenicity via a recurrent neural network[20]. 55-mer neoantigen predicted via Neo-intline was proved both in vivo and in vitro to induce CD4^+^ T cell immune response[21]. Current predictive models primarily rely on the static assessments of epitope-MHC-II interactions via the epitope/MHC-II sequences along with experimental data. However, the predictive tools for MHC-II epitopes are currently less precise, and the predicted epitopes lack strong enough immunogenicity in vivo. One possible reason is that using the sequence-based information of MHC-II epitopes combined with either experimentally obtained binding affinity or immunogenicity data to train artificial-neural-network-based models is insufficient to cover the critical factors for identifying antigenic MHC-II epitopes. The number of reported MHC-II neoantigens is even smaller[22].

In this paper, we comprehensively investigated the three-dimensional (3D) structures of 133 epitope-MHC-II complexes (pMHC-IIs) and discovered the critical structural features of MHC-II epitopes that can be expanded on MHC-II neoantigens. With the hypothesis that 3D structural features are essential factors in MHC-II epitope/neoantigen identification, we present EpiMII (EpiMPNN-MHCII), a graph-neural-network (GNN)-based model leveraging the message-passing neural network (MPNN), for MHC-II epitope design. The function is to optimize the sequences of reported MHC-II epitopes or neoantigens to achieve parallel or even higher CD4^+^ T cell immunogenicity in vivo. Rather than relying only on epitope sequences, EpiMII is trained on our novelly built 3D structure-based MHC-II epitope dataset. Such data, the first 3D dataset of MHC-II epitopes, could enable new prediction or design models to combine with multiple features, including MHC-II binding affinity and T cell immune response. EpiMII achieved 66.7% sequence recovery for the test set of our dataset and 79.0% sequence recovery for the 3D structures of the existing 103 MHC-II epitopes. Taking hepatocellular carcinoma as case studies, we examined our model’s utilization of designing neoantigens to serve as an agent of T-cell therapeutic vaccine. In-vitro assessments of IFN-γ, TNF-α and IL-4, and in-vivo treatments of tumor mouse model were successfully conducted. Here, we proposed that EpiMII is the first design model for MHC-II epitopes/neoantigens and that its robust performance gains are achieved by combining the novel 3D structure-based training data with a GNN model, further confirming our hypothesis.

## Results

### A Novel 3D MHC-II Epitope Dataset

Currently, only 133 available epitopes from Protein Data Bank (PDB) (organized as a 3D dataset) that cover 35 types of antigens have ready-to-download 3D structures, which are not enough to train a model (Figure 1A). Structural alignment of 133 crystallized MHC-II-epitope complexes (pMHC-IIs) revealed well-aligned central regions of epitopes, with more flexibility observed in the N- and C-terminus. Importantly, residues at positions 1-10 of epitopes were identified as crucial for MHC-II binding. The conserved and non-conserved residues on the MHC-II binding grooves play key roles in binding affinity and specificity. While no distinct sequence pattern emerged from the analysis of antigenic epitopes, structural overlap in epitope backbones was evident. Detailed findings, including structural data, motif conservation, and key residue interactions, are further elaborated in the supplementary information (Figures S1 to S5, and Table S1). These insights underscore the importance of 3D structural features in designing epitopes that enhance T-cell activation while maintaining similarity to the original antigen, ensuring effective immune responses.

**Figure 1.**
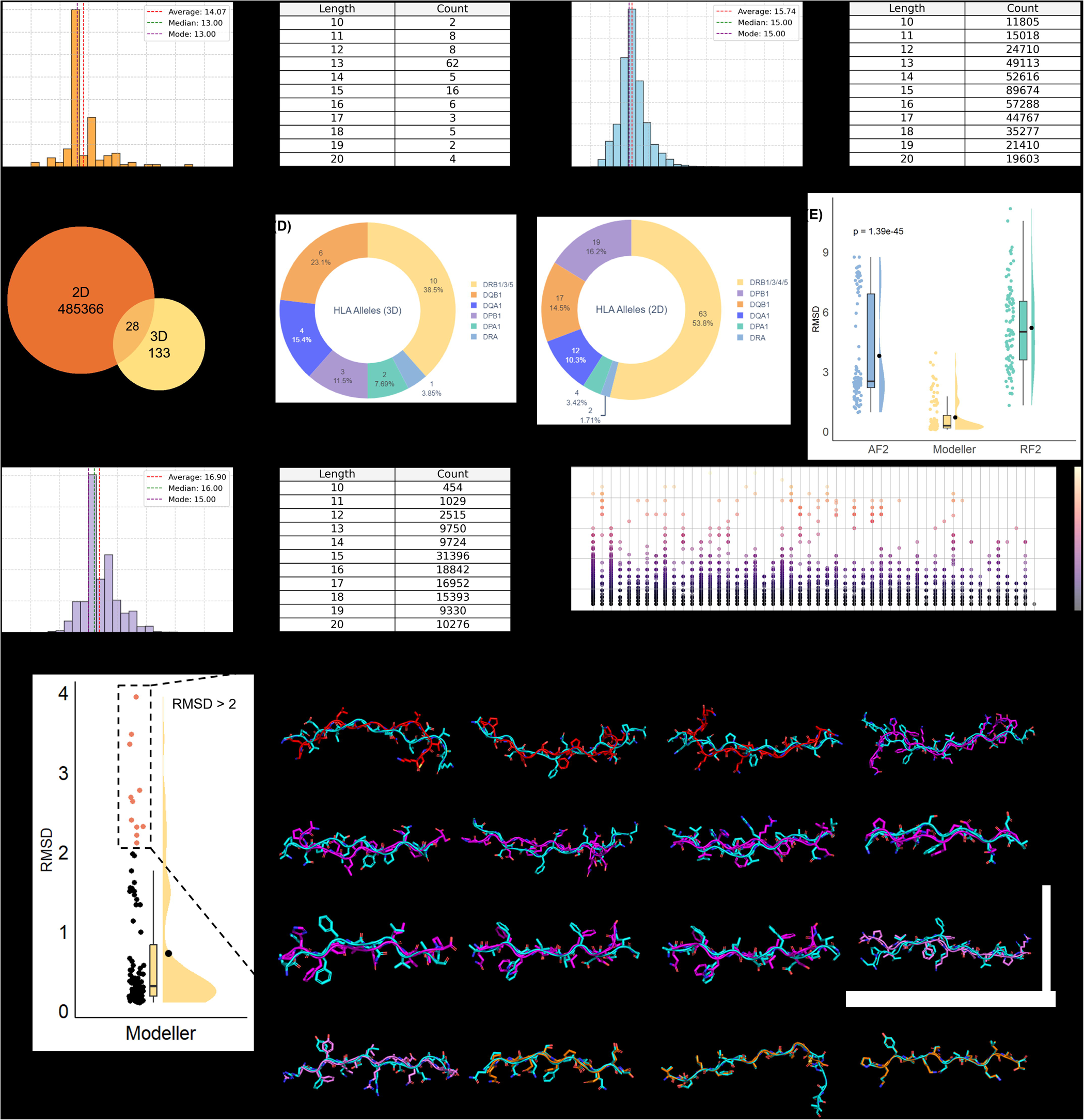
The comparison of the 3D dataset, 2D dataset and our dataset. (A) The distribution and counts of co-crystalized epitopes’ sequence lengths of the 3D dataset collected from PDB. (B) The distribution and counts of epitopes’ sequence lengths of the 2D dataset downloaded from the Immune Epitope Database. (C) The total number of epitopes in the 2D and 3D datasets. Twenty-eight co-crystalized epitopes in the 3D dataset are also included in the 2D dataset. (D) The types of HLA class II alleles are covered in the 3D and 2D datasets. For the 2D dataset, DRB has a total of 63 types covered in it, which include DRB1 (53), DRB3 (4), DRB4 (3), and DRB5 (3). (E) The homology modeling performance of three techniques, AlphaFold2 (AF2), Modeller, and RosettaFold2 (RF2), using 103 co-crystalized epitopes without modifications. RMSD values are calculated between the modeled epitopes and the co-crystalized epitopes. RMSD: AF2 3.80 Å, RF2 5.21 Å, Modeller 0.71 Å. One-way ANOVA result: p = 1.39e-45. (F) The distribution and counts of modeled epitopes’ sequence lengths in our dataset, which has 142,934 modeled epitopes. The average length is 16.9, and the median length is 16.0. The right panel shows the scatter plot of the sequence similarities of 142,934 epitopes compared to their templates. 55 templates are listed in x-axis. (G) The detailed modeling performance of Modeller. The left panel shows the RMSD calculated between co-crystalized and Modeller-modeled epitopes. The mean RMSD is 0.71 Å, and the median RMSD is 0.30 Å. The right panel shows the pairs of modeled/co-crystalized epitopes with different ranges of RMSD values. 11 pairs in total have RMSD > 2 Å (magenta and red). Three pairs have RMSD > 3 Å (red). Example pairs with RMSD < 2 Å and > 1 Å (violet). Example pairs with RMSD < 1 Å (orange).

Hence, we used homology modeling techniques to convert epitope sequences into 3D structures. We obtained 485,366 epitope sequences from IEDB and organized a sequence-based dataset covering ∼20,031 types of antigens, named the 2D dataset, to distinguish it from the 3D dataset (Figure 1B). Twenty-eight co-crystalized epitopes in the 3D dataset are also included in the 2D dataset (Figure 1C). The 2D dataset covers more diverse HLA class II allele types than the 3D dataset (Figure 1D). We employed 103 co-crystalized epitopes without modifications from the 3D dataset to test the modeling performance of three tools, RosettaFold2 (RF2), AlphaFold2 (AF2), and Modeller. The root-mean-square deviations (RMSDs) were calculated between the modeled epitopes and the co-crystalized epitopes. The results showed that epitopes modeled by Modeller have significantly lower RMSDs when compared to those modeled by AF2 and RF2 (Figure 1E, Table S2). The mean RMSD of Modeller-Modeled epitopes is 0.71 Å, and the median RMSD is 0.30 Å, which indicates an ideal homology modeling performance. Modeled epitopes with RMSDs smaller than 2 Å show excellent structural alignments on both backbone and side chain (Figure 1G). Only 11 modeled epitopes out of 103 have RMSDs larger than 2 Å but still smaller than 4 Å (Figure 1G). Thus, we determined to use Modeller to convert epitope sequences to 3D structures.

To improve the dataset quality, we performed the sequence alignment of the epitopes in the 2D dataset using 55 non-repeated, co-crystalized epitopes without modifications from the 3D dataset as templates and calculated sequence similarities. Each sequence of all 2D epitopes will be matched with one sequence among the 55 templates with the highest sequence similarity. We created three temporary datasets by selecting epitopes with the highest sequence similarities larger than 30%, 28%, or 25% and further filtered out the redundant ones. This step evaluated which sequence similarity cutoff would provide good homology models for our situation. Three datasets of modeled epitopes were used to train our model to identify which sequence similarity of epitopes can achieve better training, validation and test performance. The training results of the three datasets showed that the dataset with sequence similarity larger than 25% had the lowest validation perplexity, highest validation accuracy, and less overfitting compared to the other two datasets (Figures S6). We randomly selected 100 epitopes from the 2D dataset that were not included in the current dataset as a test set to calculate the sequence recovery. The group with > 25% sequence similarity had the highest mean and median sequence recovery among the three groups (Figure S6). We finally chose the dataset with > 25% sequence similarity. This new dataset contains 142,934 non-redundant epitopes with both sequences and Modeller-modeled 3D structures. The modeled structures have a greater than 25% sequence similarity compared with co-crystalized epitopes, which are most suitable for training a protein design model. This novel dataset will be used further for model training.

### EpiMII: A Functional Model for MHC-II Epitope and Neoantigen Design

EpiMPNN-MHCII (EpiMII) is constructed based on the message-passing graph neural network (MPNN). MPNN propagates node features by exchanging information between neighboring nodes[23]. EpiMII takes the epitopes’ structures and sequences in our dataset as inputs and learns from the 3D coordinates and amino acid composition to generate a virtual 3D shape of the epitope backbone. Then, based on the original residue composition, the residues of the epitope sequence are iteratively designed and optimized, generating multiple sequences with different sequence recoveries (Figure 2A, 2B). To ensure that the model changed only a small number of residues, we used three masking strategies to rescue the epitope with low sequence recovery to achieve different purposes (Figure 2C). 1) Fixing conserved residues to design non-conserved residues is to design hypervariable region with the potential to change the epitope binding specificity. 2) Fixing non-conserved residues to design conserved residues is to explore the alternative core binding residues out of the inherent motifs of antigenic epitopes to potentially adjust immunogenicity. 3) Masking important residues can customize the design positions based on the reported information derived from crystallography or literature. The modeling performance is measured using the test set, which contains 14,294 modeled epitopes split in our dataset (Figure 2D). The mean sequence recovery is 66.7%. Among them, only 14.34% of the epitopes in the test set have sequence recoveries lower than 0.5, indicating that most of the designed sequences have higher than 50% sequence identity compared to the original input sequence (Figure 2D).

**Figure 2.**
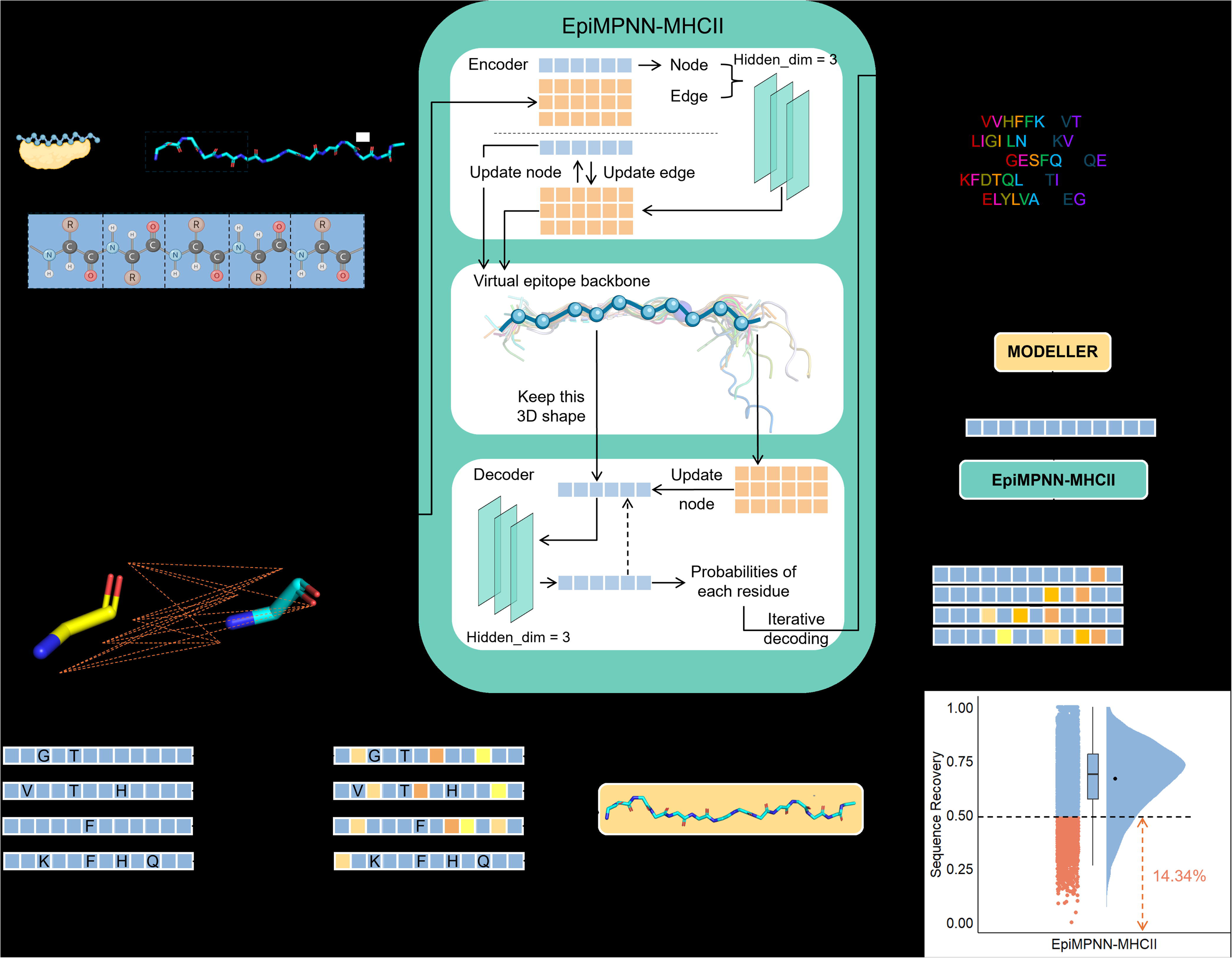
An overview of EpiMII (EpiMPNN-MHCII). (A) EpiMII takes modeled 3D epitopes as input, extracts the 3D coordinates and amino acid composition, and then calculates the distances between the same or different types of atoms, including N, Cα, C, O, and virtual Cβ. The distances are input as edges. The node is empty. The node and edges are updated through three hidden layers in the encoder layer. The updated node and edges form a virtual 3D epitope backbone shape in the latent space. In the decoder layer, nodes are updated based on the original amino acid composition via three hidden layers with a prerequisite of keeping the 3D shape. Each residue will be calculated using probabilities, and a weighted probability for the designed sequence will be generated. During the iterative decoding process, sequences are optimized to achieve better probabilities. EpiMII outputs several 2D sequences with different sequence recoveries. (B) The example of model utilization. The neoantigen will be matched with a 3D co-crystalized epitope as a template and go through homology modeling via Modeller. EpiMII will input the modeled 3D neoantigen to design its sequence, generating designed 2D sequences in the same length with different sequence recoveries. (C) Three mask strategies to improve the low sequence recoveries of the designed peptides, including masking (non-)conserved residues and masking important interacting residues. (D) The model performance measured using a test set (14294 modeled epitopes). The mean sequence recovery is 0.667, and the median sequence recovery is 0.688. Red dots represent epitopes with lower than 50% sequence recoveries that only account for 14.34%.

We also measured the influence of backbone noise ranging from 0 to 0.9 Å on the training and validation results (Figure 3A, 3B). Both training and validation perplexity are increasing while the training and validation accuracy are decreasing when adding up the backbone noise during model training. Given a backbone noise of more than 0 Å, the model will reach a low accuracy of 0.5. Meanwhile, the model is overfitting regardless of backbone noise (Figure 3A, 3B). Using 103 modeled epitopes to test the sequence recoveries of all models trained with different backbone noise, the model with no backbone noise during the training process demonstrated the highest sequence recovery, 0.78 (Figure 3C). To overcome the model’s overfitting, we chose to stop early during the training process at epoch 50, step 47050, as our final model (Figure 3D, 3E). We then test the sequence recovery by employing the 103 either co-crystalized or modeled epitopes to evaluate the model performance between EpiMII and another MPNN-based protein design model, ProteinMPNN. EpiMII was tested with co-crystalized epitopes (EpiMII_C) and modeled epitopes (EpiMII_M), while four ProteinMPNN models using different model weights were tested with co-crystalized epitopes. With the prerequisite of keeping the test input format the same as the training input format for each model, we compared the performance between ProteinMPNN models trained with co-crystalized epitopes and EpiMII_M. EpiMII_M had significantly higher sequence recovery (Figure 3F, Table S3). However, although the Modeller-modeled epitopes had minor RMSD values with the corresponding co-crystalized epitopes as we mentioned previously, EpiMII_C had much lower sequence recovery than EpiMII_M. The reason might be the input format of EpiMII training process was the Modeller-modeled epitope rather than X-ray crystallography structures. The text format of the two pdb structures illustrated that the differences lay in two areas: 1) The x, y, and z coordinates indicated two different spatial places. The most possible reason was that before Modeller starts modeling, the epitope sequence would be first aligned with a co-crystalized template, causing the 3D coordinates to shift, but the overall 3D shape was unchanged. This explained why modeled epitopes had different positions as co-crystalized epitopes. 2) The temperature factor (B-factor) was different, which described the mean displacement of the atom from an average position. The higher the B-factor, the higher the region’s flexibility is, indicating that this region has a low-resolution structure. The Modeller-modeled epitopes had slightly different B-factors compared to the co-crystalized epitopes, which might be one of the explanations that made the results of EpiMII_M and EpiMII_C different. Nevertheless, the Modeller-modeled epitopes can still be regarded as low-resolution epitopes with a highly similar 3D backbone shape to the high-resolution co-crystallized epitopes (Figure 1G). The modeled epitopes in our novel dataset were proven to train EpiMII well (Figures 3D, 3E, 3F).

**Figure 3.**
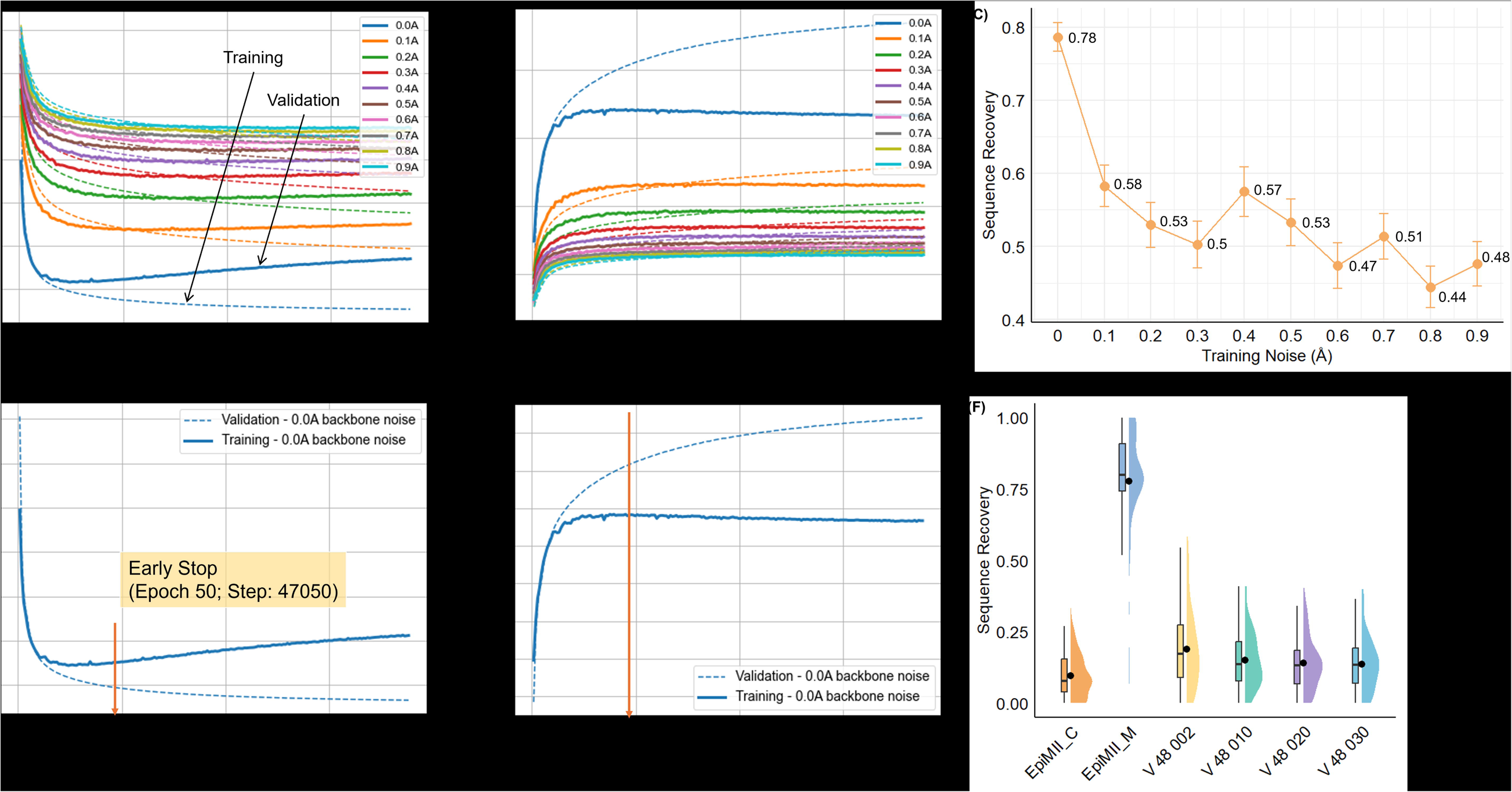
Model performance and evaluation. The training and validation performance of EpiMII with different backbone noises, ranging from 0 to 0.9 Å, is shown in (A) perplexity and (B) accuracy. Training results are shown as dash lines, while validation results are shown as solid lines. When the backbone noise is 0 Å, EpiMII has the lowest training perplexity and highest training accuracy. But all models are overfitting. (C) The output sequence recovery of models with different training backbone noises using the last epoch as model weight and 103 modeled epitopes as input. (D) and (E) show the training and validation perplexity and accuracy of EpiMII with backbone noise of 0 Angstrom, respectively. The red solid line showed the early stop at epoch 50, step 47050, to overcome overfitting, our final model weight. (F) The model performance is measured by testing the sequence recovery among EpiMII and ProteinMPNN. EpiMII_C means our model uses 103 co-crystalized epitopes to test sequence recovery. EpiMII_M means our model uses 103 modeled epitopes to test sequence recovery. ProteinMPNN uses 103 co-crystalized epitopes to test sequence recovery. The black dots mean the average sequence recovery in each group: EpiMII_M 0.78, V_48_002 0.19, V_48_010 0.15, V_48_020 0.14, V_48_030 0.14, EpiMII_C 0.10.

### Designing a Reported Neoantigen for Hepatocellular Carcinoma Using EpiMII

Hepatocellular carcinoma (HCC), constituting more than 90% of the primary tumor of the liver, is one of the most common causes of cancer worldwide[24]. The five-year survival of patients with HCC is only 18%[25, 26]. As a real-world case study, we applied EpiMII to design AYHASKYEFLANLHIT, a reported HCC neoantigen which has a single point mutation of the wild type protein encoded by the gene DUSP5. The neoantigen was selected from the Neoantigen database (Neodb: https://liuxslab.com/Neodb/), not included in our dataset, and was reported to bind to HLA-DRB10101 encoded MHC-II. When designing the original neoantigen AYHASKYEFLANLHIT (here, we named it as ‘Origin’), EpiMII finally generated 21 designed sequences with different sequence recoveries using no masking strategy, mask conserved residues, mask non-conserved residues, and mask important interacting residue strategies (Figure 4A, Table S4). We then predicted the 3D structures of all designed sequences via the AF2-multimer and AlphaFold3 (AF3) to select the ones with similar 3D shapes as we previously identified. The ptm and iptm scores have little differences for all six candidate-G (G-domain of DRA0101-DRB10101) complexes (Figure 4A, Table S4). C-ImmSim was also applied to all designed sequences to computationally assess their ability to induce the CD4^+^ T cell immune response[27]. Finally, five designed sequences that cover different masking strategies and sequence recoveries were selected for in-vitro and in-vivo experimental validations (Figure 4A, Table S4).

**Figure 4.**
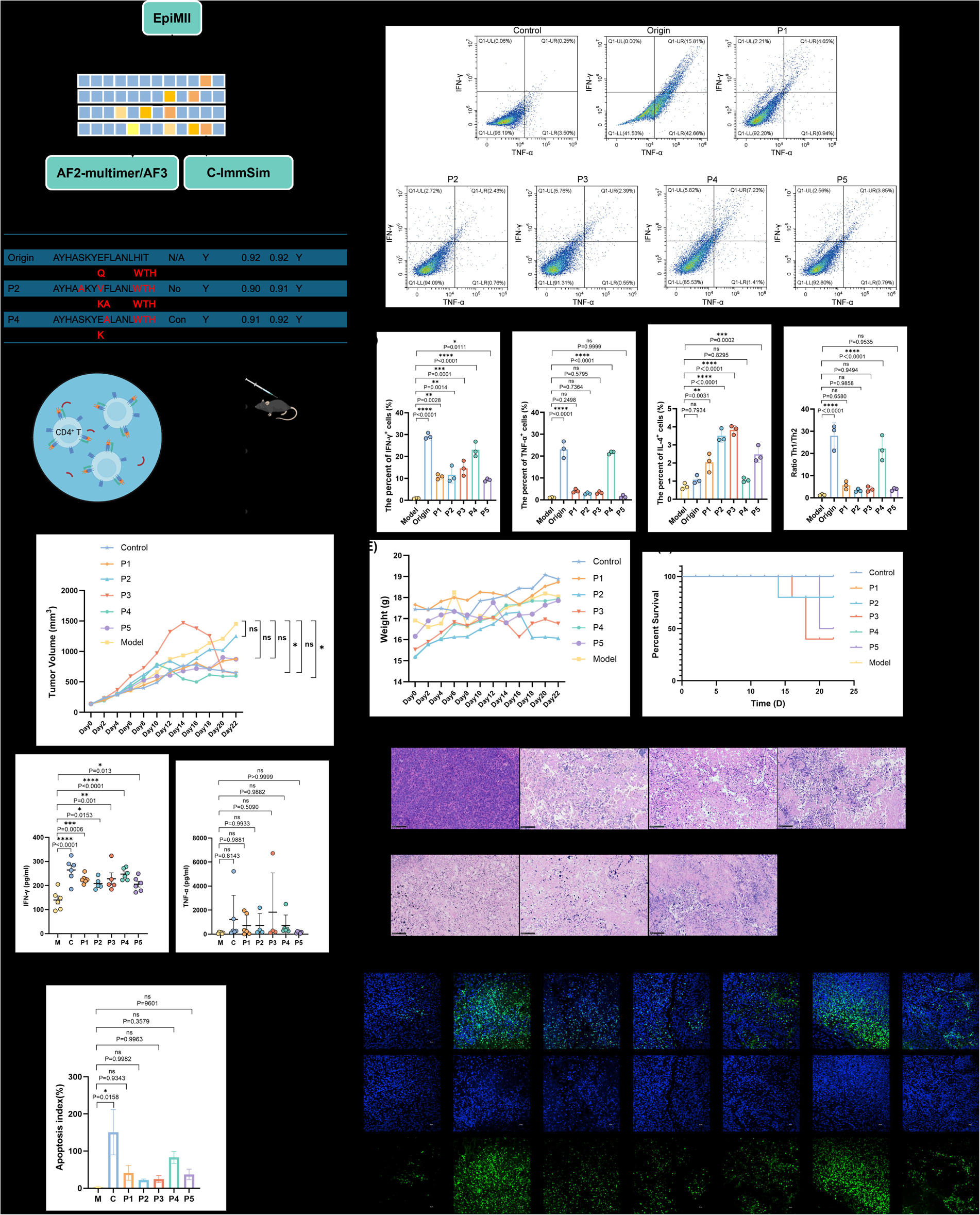
In-vitro and in-vivo experimental validations of designed HCC neoantigen candidates. (A) An overview of the whole process of candidate selection and experimental validations. (B) Trends of IFN-γ and TNF-α contents in each group. IFN-γ detected by PE, TNF-α detected by PC7. (C) IFN-γ, TNF-α, IL-4 contents (Figure S7), and Th1/Th2 ratio in cells of each group. (D) Comparison of tumor volume size in mice in 0-22 days. C vs M: P = 0.0376 (*). P1 vs M: P = 0.1221 (ns). P2 vs M: P = 0.9961 (ns). P3 vs M: P = 0.9105 (ns). P4 vs M: P = 0.0189 (*). P5 vs M: P = 0.1560 (ns). (E) Body weight of mice from 0-22 days. (F) Survival status of mice in 0-22 days. The survival rate of P2 and P4 mice was 80%, P5 50%, P3 40%, and the other groups 100%. (G) Comparison of IFN-γ and TNF-α levels in mouse serum. (H) HE staining of tumor sections in each group (Original magnification: ×40). The tumor cells in the Model group had normal volume, appearing round or oval, clear cytoplasm staining, clear nucleoli and nuclear membrane staining, and clear staining resolution. The tumor cells of the Control, P1-5 groups showed significant shrinkage, low chromatin clarity, and apparent shrinkage of the nucleolus and nuclear membrane. The nuclear envelope details are blurred. The coloring hierarchy and contrast are not clear enough. (I) Quantitative analysis of and TUNEL. (J) TUNEL staining of tumor sections in each group (400×). DAPI stained nuclei were blue at 405 excitation wavelength, TUNEL kit was labeled with green fluorescein, and positive apoptotic nuclei were green with 488 excitation wavelength.

For in-vitro validation, human peripheral blood mononuclear cells (PBMCs) were obtained from a healthy donor with the DRB10101 genotype. The immune response was stimulated by culturing PBMCs with HLA-DRA0101 and HLA-DRB10101 genotypes in vitro. The CD3^+^CD4^+^ T cells were first screened out by FITC and PerCP/Cyanine 5.5 labeling. Then, the intracellular factor fluorescence staining of PE, PC-7, and APC was detected. The results showed that Origin and P4 groups had the most potent ability to activate T cells, and the increasing trends of IFN-γ and TNF-α contents reached 15.81% and 7.23%, respectively (Figure 4B). P1 and P5 reached 4.65% and 3.85%, respectively (Figure 4B). The lowest level of T cell activation was found in P2 and P3, only 2.43% and 2.39%, respectively (Figure 4B). P4 candidate has the most significant immune activation on human CD4^+^ T cells besides the Origin neoantigen. The secretion of IL-4 was almost negatively correlated with that of IFN-γ: Origin and P4 secreted the least IL-4 compared to other groups, while P2, P3, and P5 had more IL-4 secretion compared with Origin, P1, and P4 (Figure 4C, Figures S7). IFN-γ is the surface marker of Th1 cells, and IL-4 is the surface marker of Th2 cells. According to the secretion of IFN-γ and IL-4, Th1/Th2 can be obtained. In all groups, only Origin and P4 showed excellent significant differences (Figure 4C). Thus, P4 has a good stimulatory effect on CD4^+^ T cells, reflected explicitly in the secretion of high levels of IFN-γ and TNF-α, and can effectively stimulate CD4^+^ T cells to differentiate into Th1 cells. It is preliminarily inferred that P4 can effectively activate CD4^+^ T cells, which animal experiments will further verify.

In-vivo validation was conducted on C57BL/6 mice. Mice treated with Origin neoantigen were set as the positive control group (Control). After 0-22 days of epitope treatments, the peptides with the most significant tumor inhibition effect are Control and P4 groups with P < 0.5, as shown in Figure 4D, Figure S8. P3 also showed an inhibitory effect in the later stage, but this is likely due to the error caused by the increased mortality of this group of mice in the later stage. Compared with the Model group, P1 and P5 also slowed down tumor volume growth (Figure 4D, Figure S8). The mice in all treatment groups had no significant weight loss in the past 22 days (Figure 4E). The survival rate is shown in Figure 4F. The levels of IFN-γ and TNF-α in mouse serum were measured by ELISA. All groups (C, P1-P5) have significantly higher IFN-γ concentrations than the Model group. Among them, the IFN-γ levels in the Control and P4 groups are considerably higher than others (P < 0.0001) (Figure 4G). The differences in the secretion of TNF-α are mainly reflected at the individual level: The TNF-α levels of some individuals in the Control and P3 groups have increased significantly; some individuals in P1, P2, and P4 have also increased to some extent, with P5 being the lowest (Figure 4G). The tumor cells on the HE-stained sections of the treatment groups (C, P1-5) showed significant shrinkage, low chromatin clarity, and apparent shrinkage of the nucleolus and nuclear membrane compared to the tumor cells in the Model group. Control and P4 are the most apparent (Figure 4H). TUNEL staining is consistent with the results of HE staining, showing that the apoptosis of control and P4 tumor cells was the most serious, and there was no significant difference in other groups (Figure 4I, 4J). This further confirmed the significant tumor-inhibitory effects of P4 compared to mice without any neoantigen treatment and the parallel effects compared to the positive control.

## Discussion

This study aimed to investigate the critical roles and potential utilizations of 3D structural features of antigenic MHC-II epitopes during identification. Through the thorough structure- and sequence-based analysis of MHC-II and epitopes, we created the first novel 3D MHC-II epitope dataset using homology modeling techniques to overcome the structure data insufficiency, making the structural features available in the deep learning area. Our model EpiMII achieved a 4.2x improvement in the sequence recovery of the existing 103 MHC-II epitopes derived from PDB. Using a reported HCC neoantigen as the case study, EpiMII successfully generated five neoantigen candidates demonstrating their ability to activate CD4^+^ T cells in vitro. One candidate, P4, outperformed the others regarding parallel immunogenicity and tumor shrinkage compared with the reported neoantigen in vivo.

These results indicate that the GNN-based model, EpiMII, can learn the relationship between the MHC-II epitope sequences and their 3D structural information to generate sequences that potentially keep the same function. Combining the experimental results with model outputs, we found that not the sequence with as minor changed residues as possible could, or say, the higher sequence recovery the epitope has, achieve better immunogenicity. The position of the changing residue matters the most, as shown in Figure 4A. P4 has a lower sequence recovery and more changed residues than P5, while P4 performed better than P5. We generated P4 using the ‘masking all conserved residues strategy’ in EpiMII by masking the residues at positions 1-6, 9, and 10 (Table S4). Compared to ‘no mask residues’ and ‘mask non-conserved residues’ strategies, we speculate that masking conserved residues can keep the most critical interactions unchanged between MHC-II and epitope and keep the random-coil shape of the epitope backbone at the same time. This possibly guarantees that the designed epitope will target the same site. However, the conserved 3D shape cannot be captured by AF2 or AF3 since the predicted structures have almost the same iptm and ptm scores as the input neoantigen (Figure 4A). The results of MHC-II epitope analysis align with the reverse-binding mode of DP-epitope complexes identified by Racle et al. (2023)[15]. Still, our dataset did not show MHC-II binding motifs similar to those discovered by MoDec. This discrepancy may be due to differences in the types of antigens covered in the datasets and the strategies for finding the motif. We tried identifying the residue frequencies at every position for all epitopes that bind to the same MHC-II types. Rachel et al. discovered several binding motifs shared by most of the epitopes in the same group[15]. The results do not conflict.

Our design model applies not only to MHC-II epitopes, but its application also extends to cancer immunotherapy, particularly in therapeutic T cell cancer vaccine. EpiMII makes it possible to rescue or improve the epitopes that triggered insufficient T cell immune response by optimizing specific residues on the epitope’s sequence to adjust immunogenicity and respond to the same target, for example, the same tumor site. Utilizing the conserved and non-conserved residues discovered in 66 types of MHC-II, EpiMII may increase the number of neoantigen candidates targeting the same site with different binding specificities to advance MHC-II-neoantigen-based cancer immunotherapy from the personal level to the population level. Meanwhile, EpiMII is much more efficient in lead identification than the random mutation scheme. For example, P5 was designed with one residue changed under the masking strategy, and the immunogenicity was maintained. The random mutation scheme that only changes one residue requires up to 19^*N^ mutations to achieve the same effect (N is the number of non-conserved residues). Not to mention multi-site mutations, the number of random mutation experiments increases exponentially (20^*N^), while EpiMII can efficiently complete it with just one design, such as P1-4, showing great potential to promote the MHC-II neoantigen identification process.

This study is limited by insufficient existing co-crystalized MHC-II epitopes accessed from PDB, which restricts us from collecting more sequence data with sequence similarity larger than 25% to generate our 3D dataset. Additionally, more case studies covering different diseases, like different tumor models, could provide more evidence on the better selection routine of the epitope candidates output by EpiMII. This study proved that MHC-II conserved residues determined the important binding positions on epitope sequences (eight conserved positions). Future studies can explore more on that and can further investigate whether the non-conserved residues on the MHC-II G-domain are critical for epitope binding specificities, for example, whether EpiMII can optimize specific residues that interact with MHC-II non-conserved residues to change epitope binding from DR to DQ with the prerequisite of targeting the same site. Besides, further utilization of EpiMII can be designing tumor associate antigen to see if can generate ‘neoantigen candidate’ that can mimic the tumor specific antigen and achieve higher CD4^+^ T cell immunogenicity.

In conclusion, this study highlights the essential role of 3D structural features in the MHC-II epitope identification process. It provides a novel 3D MHC-II epitope dataset that can train deep-learning models. This overcomes constraints posed by the limited 3D epitope data and addresses the epitope candidate’s lack of immunogenicity in vivo. EpiMII paved a new direction for designing MHC-II epitopes and could facilitate neoantigen identification, providing essential insights for precise epitope design and cancer vaccine development.

## Methods

### HLA class II alleles selection

We initially figured out the frequencies of HLA class II alleles (http://pypop.org/popdata/2008/byfreq-DP.php.html), which derived from a meta-analytic review of 497 population studies and represent approximately 66,800 individuals throughout the world as shown in Table S5[28]. Then, we calculated the HLA class II alleles population frequencies using Hardy-Weinberg proportions for genotypes. The 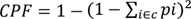, where pi is the population frequency of the i^th^ alleles within a subset of HLA-DR, HLA-DP, or HLA-DQ. The top 21 HLA-DRB1 alleles cover more than 95.54% (from DRB1*1501 to DRB1*1601), and the top 31 cover more than 99.14% of the population. The top 4 HLA-DQA1 alleles cover more than 94.9% (from DQA1*0501 to DQA1*0101), and the top 6 alleles cover more than 99.55% of the population. The top 7 HLA-DQB1 alleles cover more than 94.8% (from DQB1*0301 to DQB1*0402), and the top 16 alleles cover more than 98.5%. The top 2 HLA-DPA1 alleles cover more than 95.37%, and the top 3 alleles cover more than 99.7% of the population. The top 6 HLA-DPB1 alleles cover more than 96.0% (from DPB1*0401 to DPB1*0301), and the top 9 alleles cover more than 99.0% of the population. We chose the HLA class II alleles that cover approximately more than 99% of the global population worldwide as our samples to analyze the different types of MHC-II (plus the HLA-DRA allele), which in total are 66 alleles, including DRA (1), DRB1 (31), DQA1 (6), DQB1 (16), DPA1 (3), and DPB1 (9).

Then, we collected the 66 HLA class II alleles’ sequences from HLA Nomenclature (http://hla.alleles.org/alleles/text_index.html), accessed October 9, 2022.

### G-domain lengths determination

G-domain is the International ImMunoGeneTics Information System (IMGT) (http://www.imgt.org) unique numbering for the MHC binding grooves. For MHC-II, it is the peptide binding domain composed of the α1 and β1 regions of the chains. By searching IMGT using ‘PDB’ as the IMGT entry type and ‘MH2’ as the IMGT receptor type, we found 205 entries containing MHC-II protein. We then analyzed 11 IMGT entries covering different kinds of MHC-II, including DR, DQ, and DP. We found that the lengths of the G-domain vary not only from the alpha and beta chains but also from different types of MHC-II, as shown in Table S6. For further study, we established the general length of the G-domain, with G-alpha spanning from sequence region 1 to 84 and G-beta from sequence region 1 to 93.

### Pairwise Euclidean distances and pairwise correlation

After we obtained the protein sequences of 66 HLA class II alleles, we selected 26 mutation points for the G-alpha sequence and 50 for the G-beta sequence to represent the MHC-II binding groove and then calculated the 10 Kidera Factors for them. The 10 Kidera Factors describe multiple properties of the protein sequence[29]. The 10 Kidera Factors are including: KF1: Helix/bend preference; KF2: Side-chain size; KF3: Extended structure preference; KF4: Hydrophobicity; KF5: Double-bend preference; KF6: Partial specific volume; KF7: Flat extended preference; KF8: Occurrence in alpha region; KF9: pK-C; KF10: Surrounding hydrophobicity[29]. We calculated the Euclidean distance of the 10 Kidera Factors for 66 G-alpha domains and 66 G-beta domains and finally generated a 132*132 correlation matrix, as shown in Figure S1A.

We then searched for the reported 15-mer epitopes bound to different MHC-II types on the Immune Epitope Database (IEDB) (https://www.iedb.org/). We chose ‘Host’ as Human, and because in our 66 HLA alleles, the alleles encoded for the alpha chain are the minority compared to the other encoded for the beta chain, we selected the ‘MHC Restriction’ to be the DxB alleles in our 66 HLA class II alleles. The detailed numbers of selected epitopes for each DxB allele are shown in Table S7. We then measured each DxB group’s Pearson correlation and generated a 51*51 matrix, as shown in Figure S1A.

We collected the epitopes for HLA alleles with > 0.9 sequence similarities and removed the repeated epitopes because some epitopes are reported to bind to several HLA alleles. The sequence logo plots were generated for the eight HLA alleles groups. Then, we tried to discover the potential motifs in each allele group using MEME[30]. MEME is an online tool to discover novel, ungapped motifs in the sequences (https://meme-suite.org/meme/tools/meme). We chose the ‘Classic mode’ and requested ten motifs to find.

### Molecular Complex Characterizing System (MCCS) scoring technique

MCCS is a novel characterization method for protein-ligand complex, which includes scoring and docking techniques[31]. Here, we employed the scoring technique to characterize the binding features of by quantitating the pMHC-IIs energy contribution of each individual residue[31]. Our inputs were 133 pMHC-IIs derived from X-ray crystallography. Chimera (version 1.15) was first used to fix residues with an incomplete side chain in MHC-II PDB structures[32]. The uncompleted residues were revealed after Chimera scanned the complete protein structures. The Dunbrack rotamer library was used to replace the truncated side chains with a whole side chain of the same type of residue[33]. The polar hydrogens, Gasteiger charges, and Vina force field were added using VEGA. Finally, the PDB format of both MHC-II and epitope was converted to PDBQT format for further scoring function via jdock (version 2.2.3b, https://github.com/stcmz/jdock accessed on 20 March 2024).

Jdock is a core implementation of MCCS and is an extended variant of idock, which is used in scoring and docking functions[34]. It can generate a vector of residue-free energy from the conformation determined by the X-ray crystal structure in five terms: gauss1, gauss2, repulsion, hydrophobic, and h-bonding, which AutoDock Vina invented[35, 36]. The scoring results were nine binding recognition vectors: (1) Gauss (Gauss1 + Gauss2), (2) Gauss1, (3) Gauss2, (4) repulsion, (5) steric (Gauss1 + Gauss2 + repulsion), (6) hydrogen-bonding, (7) hydrophobic, (8) non-steric (hydrogen-bonding + hydrophobic) and (9) residue energy contribution. We chose the ‘score only’ mode for performing the scoring technique in which the scores of all receptor-ligand atom pairs were directly calculated and added to the overall score.

### 3D dataset

We searched the IMGT using ‘PDB’ as the IMGT entry type and ‘MH2’ as the IMGT receptor type, and we found 205 entries containing MHC-II protein. We then excluded the complexes that the bound peptide is a class II-invariant chain (CLIP), which blocks the antigen binding region of the MHC-II before the epitope comes. We finally obtained 133 pMHC-II and downloaded them from the PDB. Their epitopes were organized as a 3D dataset containing 29 modified epitopes and 23 mutant epitopes.

### 2D dataset

We searched the IEDB using ‘Human’ as Host, ’Class II’ as MHC Restriction, and ’include the positive assays’ and obtained around 500,000 epitopes with 2D sequences. After excluding the epitopes with three-letter amino acids, numbers, modified residues such as CIR, and lengths smaller than 5-mer, we finally got 485,366 2D epitopes.

After obtaining the list of 485,366 epitopes in a ‘.txt’ file, with the ID and sequence of each epitope stored on a single line and separated by commas, we used a script to convert the epitope information into individual ‘.ali’ format files for each epitope sequence. The script reads a list of sequence IDs and sequences from the input file, generating corresponding ‘.ali’ files in the output folder. Each ‘.ali’ file is named after the sequence ID and includes a formatted header along with the associated sequence. To build the 3D structure of each epitope, we needed to identify the appropriate template(s) for each target. We utilized 55 diverse 3D structures of reported epitopes as templates set to conduct sequence alignments between each target epitope and the known structures. Essentially, the script automates the alignment of target epitope sequences (stored in ‘.ali’ files) against a set of known 3D epitope structures using Modeller[37, 38]. During the sequence alignment, it classifies amino acids into specific categories and calculates sequence identity by comparing residue matches and similarities. The script reads the target sequences, loads the known structures as templates, and performs multiple sequence alignments to identify the best matching template for each epitope based on sequence identity. The results, including alignment files, the best template, and similarity score are saved, and all files are organized into a specified output directory. During this step, we created a dataset of 142,934 non-redundant epitope sequences, each sharing at least 25% sequence similarity, to build their 3D structures.

Using the alignment files, we then employed an in-house script to build homology models for each target epitope. This script automates the homology modeling process using Modeller (‘automodel’), iterating through the target epitope sequences in ‘.ali’ format, and identifying the corresponding template PDB structures from a predefined list. The script builds comparative models using the selected templates and target sequences, generating 3D structures of the epitopes. Each model is saved as a ‘.pdb’ file, and all relevant alignment and model files are organized into specified output directories.

### Homology modeling techniques

103 non-redundant sequences of the co-crystalized epitopes that bind to MHC-II were selected from the 3D dataset and employed to test the modeling performance of homology modeling techniques, including RF2, AF2, and Modeller. These epitopes have no modifications such as citrulline (CIR) and acetyl group (ACE). AF2, developed by DeepMind, is a neural network-based model that can predict protein structures with atomic accuracy and even accuracy in circumstances where no similar protein structure is available[39]. We accessed AF2 via the Center for Research Computing (CRC) at the University of Pittsburgh and predicted structures for 103 epitope sequences using an A100 GPU (80GB SXM) with ‘db_preset’ as ‘full_dbs’ and ‘model_preset’ as ‘monomer’. RF2 is another neural network-based protein structure prediction model that is more computationally efficient than AF2, and it has been reported to have parallel accuracy of AF2 on monomers[40]. We accessed RF2 on our server Hydra (Intel Core i9-13900K CPU, 24 cores, 128GB) and utilized identical 103 2D sequences. Modeller is used for homology or comparative modeling of protein 3D structures, which requires an alignment of a protein sequence to be modeled with the known related structures[37, 38]. We performed homology modeling using 55 3D co-crystalized epitopes as templates. Then, structural alignment was performed using Pymol (TM) 2.5.7 between the modeled/predicted epitopes in three groups and their corresponding co-crystalized epitopes to calculate the RMSD values. The average RMSD values out of 103 pairs were identified as the criteria for comparing the modeling performances of three techniques. We employed analysis of variance (ANOVA) to evaluate our testing results, shown in Figure 1E.

### EpiMII Architecture

The model architecture was encoder-decoder message passing neural networks[41, 42]. The overview was shown in Figure 2A. The encoder layer had three hidden dimensions to update the node and edges. The decoder layer also had three hidden dimensions to iteratively decode the amino acid compositions to generate epitope sequences.

Graph Representation (Encoder Processing). Peptides structures were depicted as graphs, where atoms served as nodes and interactions between atoms, such as bonds and distances, were depicted as edges. Each node (atom) was associated with a feature vector that encapsulated its atomic properties, including type, coordinates, charge, hydrophobicity, and any masked values. Similarly, each edge (interaction) was associated with a feature vector that encoded pairwise interactions between atoms, including bond type, distances (e.g., pairwise Euclidean distances), and potentially radial basis function (RBF) values. This initial representation converted the input structure into a graph format suitable for neural network processing.

Message Passing Layers (Encoder Processing). EpiMPNN-MHCII employed message passing neural network (MPNN) layers to propagate information between nodes (atoms) and update their representations based on interactions with neighboring nodes. This process involved iterative steps where each node aggregated information (messages) from its neighboring nodes and updated its representation accordingly.

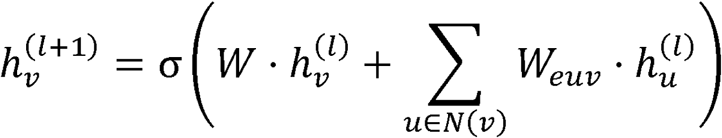

where 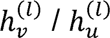 is the hidden state of node *v* / *u* at layer *l*, *σ* is an activation, *W* is a learnable weight matrix, and *W_euv_* is a learnable edge-specific weight matrix. Typically, multiple message-passing layers were utilized to capture hierarchical and spatial features of the antibody structure.

Node and Edge Update Functions (Decoder Processing). Within each message passing layer, node and edge update functions were applied to compute new node and edge representations based on aggregated information from neighboring nodes:

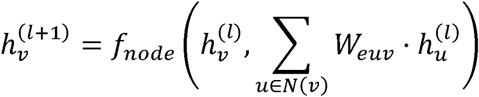

where *f_node_* is a learnable node update function.

Similarly, edge update functions process edge features and node embeddings to update edge representations, capturing spatial relationships and interactions within the antibody:

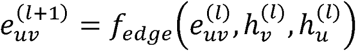

where *f_edge_* is the edge update function, 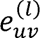 represents the updated feature of the edge (*u*, *v*) after layer *l*, 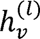 is the feature of node *v* at layer *l*, 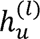 is the feature of node *u* at layer *l*.

Graph Pooling (Decoder Processing). Following multiple message-passing layers, EpiMPNN-MHCII employed graph pooling operations to aggregate node representations and generate a global representation of the entire antibody structure.

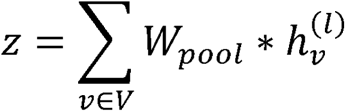

where *z* is the pooled representation, *W_pool_* is a learnable weight matrix, and *l* is the number of message-passing layers.

Training and Optimization (Decoder Processing). EpiMII was trained using supervised learning techniques on large datasets of epitope structures with known properties, such as experimental structures from databases. During training, the model learned to minimize a loss function that measured the difference between predicted and actual epitope coordinates.

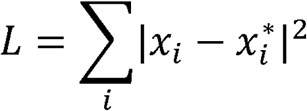

Where *L* is the loss function, *x_i_* is the predicted amino acid sequence, and *x_i_Ȫ* is the actual antibody sequence.

Optimization methods, such as gradient descent, were employed to update the model’s parameters and improve its predictive performance over iterations:

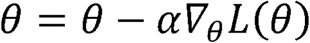

where *θ* is the set of learnable parameters, *α* is the learning rate, and *∇_θ_L* (θ) is the gradient of the loss function with respect to the parameters.

Sequence Generation (Decoder Processing). Through an autoregressive approach, EpiMII meticulously predicted the amino acid sequence x given the underlying backbone structures, encapsulating the intricate relationship between structural elements and sequence composition:

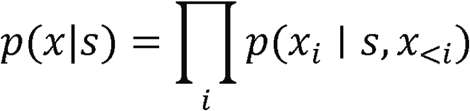

Here, *p*(*x_i_* | *s,x*_<*i*_) denoted the conditional probability of the amino acid *_i_* at decoding step *i*, and *s_<i_* = {*x*_1_ =, … *x_i−1_*} probabilities were parameterized using two primary components: an encoder that computed node and edge embeddings from structural data, and a decoder that predicted the next decoded residue autoregressively based on preceding decoded letters and structural embeddings.

### EpiMII: Model training process

Our model training process used the same model architecture and feature input as ProteinMPNN. For the training dataset, we first tried to cluster the 485,366 Modeller-modeled MHC-II epitopes by mmseqs2 with alignment mode three and single-step clustering method based on the sequence identity: 30%, 25%, 20%, 15%, 10%[43, 44]. The dataset resulted in 274,353 clusters for sequence identity 30%, and 274,063 clusters for sequence identity 25%, 20%, 15%, and 10%. The models trained on these clusters all resulted in inadequate training and validation performance, in which validation perplexities were higher than seven, and accuracies were lower than 0.40. We then chose to clean the 2D dataset based on sequence similarity that larger than: 30%, 28%, and 25%, meanwhile, made each epitope in the dataset after cleaning as one cluster. Sequence similarity was calculated as the percentage of similar residues in two sequences. The residues in two sequences that belong to the same biochemical groups, including nonpolar/hydrophobic, polar uncharged/hydrophilic, acidic/negatively charged, basic/positive charged, and others, were identified as similar residues. The dataset with larger than 30% sequence similarity had 20,022 epitopes. The dataset with larger than 28% sequence similarity had 44,019 epitopes. The dataset with a sequence similarity larger than 25% had 142,934 epitopes in total. We trained models on these three datasets and tested the sequence recovery using 103 modeled MHC-II epitopes, as shown in Figure S6. Sequence recovery is a benchmark method of evaluating protein design tools, which passes the backbone of natural protein with the known amino-acid sequence as input. It measures the identity between the predicted and true sequences to evaluate the prediction accuracy[29]. We also trained the model with 485,366 clusters (each epitope in the 2D dataset was regarded as one individual cluster) without clustering on either sequence identity or sequence similarity. Although the training and validation performance of this model were parallel to the model trained on the dataset with sequence similarity > 25%, the tested sequence recovery using 103 modeled epitopes (mean average: 0.6623, median average: 0.7308) was worse than the dataset with sequence similarity larger than 25% (mean average: 0.7865, median average: 0.8375). Thus, we determined to use 142,934 Modeller-modeled non-redundant epitopes (sequence similarity > 25%) with their 3D structures and 2D sequences as our dataset in this study.

The dataset was split into train, validation and test sets in 8:1:1. Each modeled epitope was set up as one cluster. We trained the model with the setup: ‘hidden_dim’ as 128, the number of neighbors for the sparse graph as 48, epoch as 200, ‘batch_size’ with 2048 tokens, reload data every 4 epochs, save model every 2 epochs, and the ‘gradient norm’ is -1. The dropout rate is 0.1, and the backbone noise is 0 Angstrom. We used the learning rate schedule as the ProteinMPNN paper[41]. Models were trained using automatic mixed precision. The input features of the model were embedded edges without node features, which were the Euclidean space and distances between the residues in the primary sequence space with the same chain ID indicator, ‘_A’. The distances between two residues are encoded using the Gaussian radial basis functions (RBFs) equally spaced from 0 Angstrom to 20 Angstrom between atoms, N, Cα, C, O, Cβ. The virtual Cβ coordinates were calculated as β = −0.58273431 * a + 0.56802827 * b − 0.54067466 * c + Cα − N,c = C − Cα,a = cross(b, c) which represents the ideal angle and bond length definitions.

We kept the model’s loss function and optimization section the same as ProteinMPNN: 1) The training loss was defined by: . 2000 was chosen empirically, loss (categorical cross entropy per token) and mask had shapes [batch, protein length]; 2) The optimization was using Adam with beta1 = 0.9, beta2 – 0.98, epsilon = 10-9. The models were trained on CRC of the University of Pittsburgh, using pytorch 2.0.1, cuda 11.8 with an A100 GPU, and a memory of 40GB. The training and validation losses (perplexities) converged around 150k-200k optimization steps, about 200 epochs of the training data.

### EpiMPNN-MHCII: Model utilization

We used the early stopped model weights on epoch 50, step 47050, as the final model to design the sequences of neoantigen samples. We modeled the sequences of neoantigens to the 3D structures via Modeller and input into our model. The backbone noise was set to zero, and the number of sequences to be generated was set to 16. The ‘designed_chain’ was the single chain of the neoantigen, and no chain was fixed (fixed_chains = []). The outputs were the original neoantigen sequence and 16 sequences of the designed neoantigen. Designed peptides and their sequence recoveries (seq_recovery) were recorded. As for the masking strategy, we first aligned the modeled neoantigen structure with the standard co-crystalized epitope extracted from ‘1bx2.pdb’ to number the residue positions of the neoantigen sequence. Then, the positions to be masked on the neoantigen sequence can be set up through ‘fixed_positions_dict’. Our mask strategies are shown in Figure 2C. We used ‘mask no residues’, ‘ mask conserved positions’, ‘mask non conserved positions’ and ‘mask the positions of important interacting residues’ to generate a list of sequences with different sequence recoveries. Because the masked residues did not participate in the calculation of sequence recovery, we finally selected the sequences with no more than five residues changed compared to the original neoantigen sequence for further validation.

### AlphaFold2-multimer and AlphaFold3

AF2 is a neural network-based model that can predict protein structures with atomic accuracy[39]. AF2 was accessed via the CRC at the University of Pittsburgh and ran on an A100 GPU (80GB SXM). Because AF2 is more accurate in predicting structure when we input both the sequences of epitope and MHC-II compared to the epitope sequence only, we input the sequences of the designed neoantigen and the whole MHC-II that the original neoantigen binds to make the structural prediction. The complete sequences of the alpha and beta chain of the specific type of MHC-II were collected from the HLA Nomenclature (http://hla.alleles.org/alleles/text_index.html). We then chose the epitope of 1bx2.pdb as the reference structure for the neoantigens that bind to HLA-DRB1 encoded MHC-II. We employed AF2 to predict the structures of designed neoantigens with ‘db_preset’ as ‘full_dbs’ and ‘model_preset’ as ‘multimer’. We also use AF3 to perform complex structure prediction, using the sequences of designed epitope and G-domain. AF3 was access on our own server Laker (Intel Core i9-13900K CPU, 24 cores, 128GB, A6000 GPU). We performed the structural alignment for the predicted structure of the designed epitope and the reference structure using Pymol. RMSD was also calculated. Whether the shape of the designed neoantigen is similar to the co-crystalized reference structure as well as the RMSD were recorded.

### C-ImmSim

C-ImmSim is the C-language-based immune system simulator[27]. With a given agent, this online tool can predict the immune response it may trigger in both MHC-I and MHC-II antigen presentation pathways. C-ImmSim also predicts the MHC binding. We input the designed neoantigens into it to predict the potential immune response they can trigger. C-ImmSim is available at: https://kraken.iac.rm.cnr.it/C-IMMSIM/index.php. We set the basic parameters as the default value: random seed as 12345, simulation volume as 10, simulation steps as 100, time step of injection as 1. The injecting agent is a vaccine with no LPS. The number of agents to inject is 1000. We input the designed neoantigen sequences with the FASTA format and input one sequence at a time.

### Synthesize linear peptides from the C-terminus to the N-terminus direction

The solid-phase peptide synthesis process involves sequential coupling of amino acids from the C-terminal to the N-terminal, adhering to a systematic protocol. Initially, RINK resin is soaked in dichloromethane (DCM) and washed with N,N-dimethylformamide (DMF) to prepare it for coupling reactions. Fmoc protecting groups on the resin are removed using piperidine/DMF, confirmed by ninhydrin test. Coupling of each amino acid with HOBT and DIC is conducted at 30°C, followed by acetylation and washing steps. This cycle is repeated for each subsequent amino acid until the sequence is complete. The peptide is then cleaved from the resin using a cleavage solution and purified via HPLC under specified conditions. The process ensures high purity and accuracy in peptide synthesis.

### In vitro validation of peptides

Human PBMCs were obtained from a 32-year-old healthy Chinese man with the DRB10101 genotype at Shenzhen Second People’s Hospital. Separate PBMCs and co incubate with peptides (2 µM) and add 15% fetal bovine serum and IL-2 (1.7628 µg/mL) to RPMI 1640. After 11 days, the cells were placed in RPMI 1640 supplemented with 8% human serum and without any IL-2. On the 12th day, wash the cells with RPMI 1640, dilute with 2 × 10^6^/mL, and inoculate 2 × 10^5^ cells into each well of a 24 well plate. Then add 100 µL of peptide (2 mM) to each plate and incubate the cells at 37°C for 1 hour.

Add protein transport inhibitors and incubate the cells at 37°C for an additional 4 hours. Surface labeling staining was performed using PerCP/Cyanine5.5 Anti Human CD3 Antibody, FITC Anti Human CD4 Antibody, and CD4, while intracellular factor labeling staining was performed using PE Anti Human IFN-γ Antibody, APC Anti Human IL-4 Antibody, and HU TNF PE-CY7 MAB. Flow cytometry was used for final testing.

### Mouse models

All animal experiments were approved by the Animal Ethics Committee of Zhejiang University of Technology (Approval No. IACUC-MGS20241030021). Female C57BL/6J mice (4–6 weeks old) were purchased from Hangzhou Hans Biological Technology Co., Ltd., and housed in pathogen-free facilities at the Experimental Animal Center. The mice were fed a standard diet and provided with distilled water and libitum. Sterilized cages with fresh bedding were supplied weekly. At the end of the experiment, the mice were anesthetized with sodium pentobarbital (60 mg/kg) and euthanized using sodium pentobarbital (100 mg/kg). Measures were taken to minimize animal suffering. C57BL/6J mice carrying 5 × 10 subcutaneous (s.c.) Hepa 1-6 tumor cells were randomly assigned to treatment groups (6 mice per group), with an average tumor volume of 100–150 mm³ per group. Starting from day 0, MHC-II peptides were subcutaneously injected at a dose of 20 µg twice weekly. Every other day, use a digital caliper to measure the size of the tumor and observe the weight and survival rate, and tumor volume was calculated as length × width² × 0.5. One mouse in the P2 and P3 groups died naturally during the experiment. In compliance with animal ethics rules, the data for mice with tumors that grew too fast were processed as humane endpoints before the end of the experiment.

### ELISA was used to determine the concentrations of IFN-γ and TNF-α in serum

The murine IFN-γ ELISA employs a double-antibody sandwich method to quantify IFN-γ levels in serum, plasma, or related fluids. Microplate wells are pre-coated with an anti-IFN-γ antibody, followed by the addition of standards or samples and an HRP-labeled detection antibody to form an immune complex. After washing, TMB substrate is added, producing a color change proportional to the IFN-γ concentration, which is measured at 450 nm. The concentration is determined using a standard curve, and appropriate dilutions are applied to calculate the actual sample values. The determination of TNF-α is the same.

### HE staining

Paraffin sections of tumor were deparaffinized by sequential immersion in Xylene I (8 mins), Xylene II (8 mins), Xylene III (8 mins), Absolute Ethanol I (5 mins), Absolute Ethanol II (5 mins), 85% Ethanol (5 mins), and 75% Ethanol (5 mins), followed by rinsing in tap water for 2 mins. Hematoxylin staining was performed for 6 mins, followed by differentiation in hydrochloric acid ethanol for 2 secs and rinsing with water. The sections were blued in ammonia solution for 15–30 secs and washed again with water. Eosin staining was conducted by dehydrating the sections in 95% ethanol for 1 min, followed by staining in eosin solution for 10–30 secs. For dehydration and mounting, the sections were sequentially immersed in Absolute Ethanol I (30 secs), Absolute Ethanol II (2.5 mins), Absolute Ethanol III (2.5 mins), Xylene I (2.5 mins), and Xylene II (2.5 mins) for clearing, then mounted with neutral resin. Finally, the sections were examined and analyzed under a microscope with image acquisition.

### TUNEL staining

The procedure of TUNEL staining began with deparaffinization of tissue sections by sequential immersion in Xylene I (12 mins), Xylene II (12 mins), Absolute Ethanol I (6 mins), 95% Ethanol (6 mins), and 85% Ethanol (6 mins), followed by rinsing in distilled water for 2 mins. Antigen retrieval was performed using a microwave at medium power for 8 mins with citrate antigen retrieval buffer (pH 6.0), followed by cooling at room temperature and triple rinsing with double-distilled water (5 mins each). After air-drying, a hydrophobic barrier was drawn around the tissue using a histochemical pen, leaving a 3–4 mm margin, and washed three times with pure water (5 mins each). The TUNEL working solution was prepared by mixing TdT enzyme (1 µL) with reaction buffer (50 µL), applied to the circled tissue, and incubated in a humidified chamber at 37°C for 1.5 hours (adjustable based on experimental results). The sections were washed in PBS (pH 7.4) three times (5 mins each) on a shaker, excess PBS was removed, and DAPI staining solution was added for nuclear staining under light protection at room temperature for 8 mins. After washing in PBS three more times, the sections were mounted with an anti-fluorescence quenching mounting medium. Finally, the slides were examined and imaged under a fluorescence microscope, using appropriate excitation and emission wavelengths for DAPI (UV excitation: 361–389 nm, emission: 420 nm, blue fluorescence), FITC (excitation: 465–495 nm, emission: 515–555 nm, green fluorescence).

### Statistics

We employed One-Way ANOVA data analysis to compare the modeling performance among AF2, RF2, and Modeller. The F value was 147.5248, and the factor degree of freedom was 2. The P-value was 1.39e-45. The in vitro experiments of polypeptides were repeated three times for verification. All used ordinary One-Way ANOVA with degrees of freedom of 6. The F value of IFN-γ analysis was 44.89, the F value of TNF-α analysis was 119.6, the F value of IL-4 analysis was 42.23, and the F value of Th1/Th2 ratio analysis was 33.12. In the in-vivo validation, each group’s tumor volume histograms used SEM mean statistics. The pooled line graphs were analyzed using Two-Way ANOVA data analysis with multiple comparisons between individuals. The row degree of freedom was 6, the F-value was 19.63, the column degree of freedom was 11, and the F-value was 6.26. ELISA measured the content of IFN-γ and TNF-α in serum twice with duplicate wells. Both used ordinary One-Way ANOVA. The degree of freedom of IFN-γ content was 6, and the F value was 9.833. The degree of freedom of TNF-α content was 6, and the F value was 0.8782. The apoptotic index measured by TUNEL three times was also analyzed by One-Way ANOVA with a degree of freedom of 6, and the F value was 3.789. Each group’s apoptotic index histograms used SD mean statistics.

## Author Contributions

Y.J proposed and implemented the hypothesis and methodology, conducted the computational analysis, created dataset and constructed the model. X.X conducted the in-vitro and in-vivo experimental validations. X.X, C.Y, H.T, F.Z provided guidance throughout the project. Y.J constructed the GitHub. S.Z validated the GitHub. Y.J and X.X wrote the manuscript and all authors contributed.

## Acknowledgments

The authors thank the computing resources from the Center for Research Computing (CRC) at the University of Pittsburgh.

## Data availability

The dataset (pt files), 3D dataset contains 142934 modeled MHC-II epitopes (pdb files), and all model weights are available and can be downloaded on Zenodo (https://doi.org/10.5281/zenodo.14767257). 133 co-crystalized MHC-II epitopes collected from PDB and the original 2D dataset contains around 480,000 MHC-II epitope sequences collected from IEDB can be found on the Github (https://github.com/JIY106/EpiMII.git). The raw data used in the experimental validations can be found in Zenodo (https://doi.org/10.5281/zenodo.14767107). All other data sources can be found in the Supplementary figures and tables.

## Code availability

The source code of EpiMII is freely available on GitHub under Apache-2.0 license and can be downloaded on GitHub (https://github.com/JIY106/EpiMII.git). The code for running MCCS scoring techniques can be found in Supplementary figures and tables.

