## Supplementary material for "EpiMII: Integrating Structure and Graph Neural Networks for MHC-II Epitope and Neoantigen Design": Figure S

^*^ To whom correspondence should be addressed.

**3D structures of epitopes play vital roles in binding to MHC-II.**

One of the challenges of identifying epitopes that participate in the MHC-II antigen presentation pathway is the polymorphism of HLA class II alleles. We first collected the sequences of 66 HLA class II alleles that cover around 99% of the population worldwide, including DRB1, DPB1, DQB1, DRA, DPA1, and DQA1 HLA locus. The pairwise Euclidean distances of the G-domains (MHC-II binding groove) of 66 MHC-II demonstrate high in-group similarities that appear in the same HLA locus (Figure S1A). This is consistent with the nomenclature of HLA alleles that HLA-DRB1*08 represents a group of alleles with homologous sequences that encode the same antigen[1]. Another challenge in selecting the antigenic epitope is the opening ends of the MHC-II binding groove, causing the various lengths of bound epitopes. The pairwise correlation of 15-mer epitopes bound to each type of MHC-II illustrates low in-group similarities, while eight groups show high cross-group similarities (> 0.9) (Figure S1A). It indicates that the epitopes bound to this MHC-II group have a similar amino acid frequency at each position. However, for each group, the sequence logo plots show that no specific residue has a higher frequency than others at each position (Figure S1B). We further discovered the shared motifs of each group of epitopes using MEME[2]. The most shared motif among epitopes of one group accounts for less than 6% of the total number of epitopes (Figure S1C). Thus, no common pattern has been found on the various MHC-II epitopes from the sequence perspective.

From a structural perspective, we conducted a structural alignment of 133 X-ray crystalized pMHC-IIs collected from the Protein Data Bank (PDB), using 1aqd as the reference template and computing the root-mean-square deviations (RMSDs). All three groups have low RMSDs (Figure S1D). DR-epitope complexes exhibited lower RMSDs (mean: 0.56 Å) compared to DQ/DP-epitope complexes (means: 0.80 Å) (Figure S1D). Notably, the backbone structural alignment of all co-crystallized MHC-II epitopes indicated a well-aligned middle section, while the N- and C-terminus displayed greater flexibility (Figure S1E, S1F). Two examples of the structure alignment of pMHC-IIs are shown in Figure S1G. Identical epitope sequences can either bind to different types of MHC-II or have opposite binding orientations when bound to the same MHC-II (Figure S1G). This reverse-binding mode has also been identified in DP-complexes[3]. For further analysis, we named the first highly superimposed residues as position one and numbered the other positions. The eight most overlapped epitope positions bound to all MHC-II types are identified (Figure S1F).

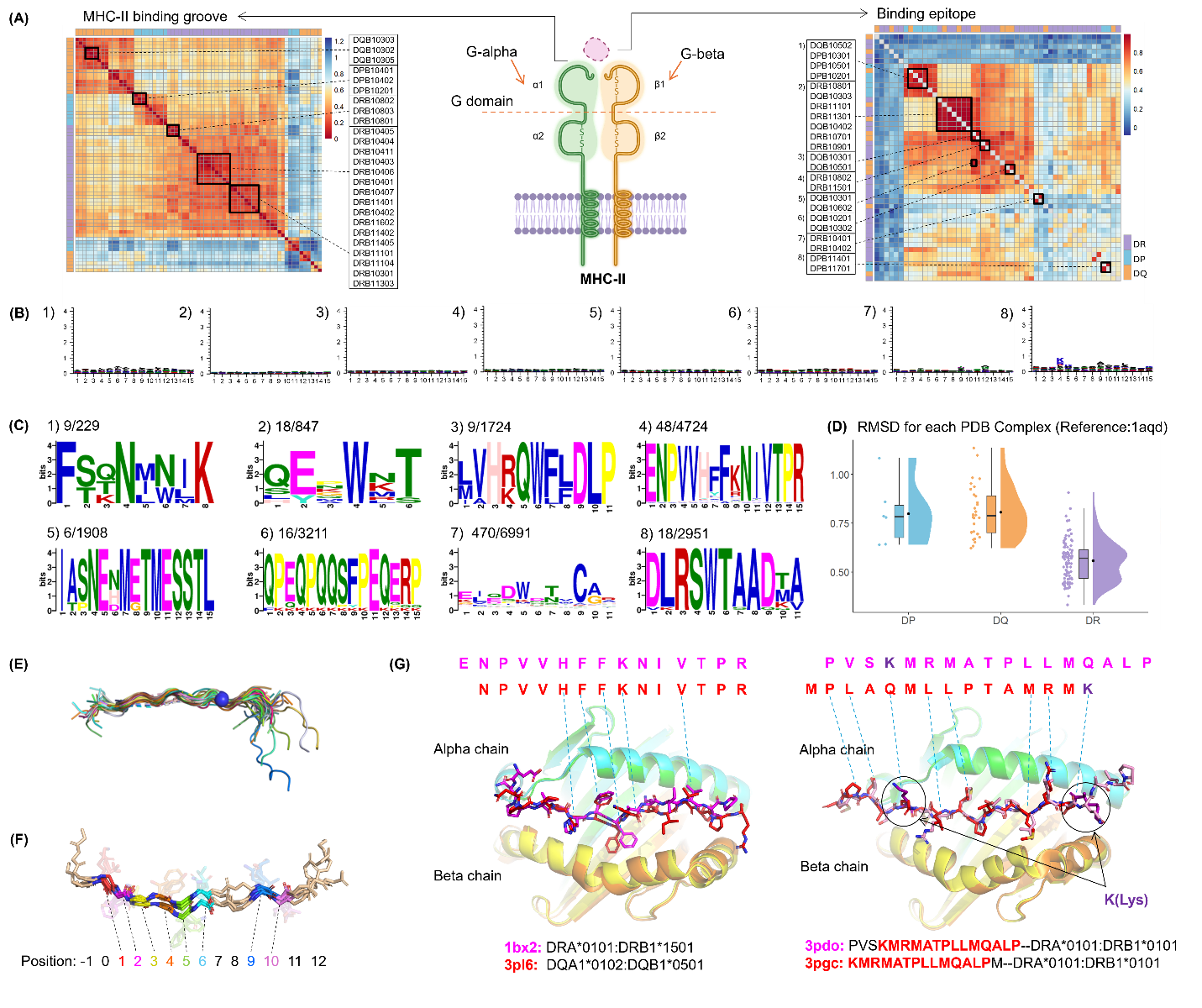

Figure S1 The overall sequence and structure analysis of MHC-II and epitopes. (A) The left panel shows the pairwise Euclidean distances of the G-domain. Some alleles with distances smaller than 0.2 are selected in black squares, indicating the more similar patterns of sequences found in these groups. The alleles' names are shown on the right. The middle part shows the overview of MHC-II-epitope complexes with a G-domain composed of the α1 and β1 regions. The right panel shows the pairwise correlation of 20 amino acids' frequency at each position of epitopes binding to specific MHC-II types. Allele types with a correlation larger than 0.9 are selected in black squares, indicating the epitopes binding to them have more similar amino acid frequencies in each position. The allele names are shown on the left. (B) Sequence logo plots of epitope sequences in the eight groups selected in (A) right panel. (C) Eight sequence logo plots of the top one shared motif in each group of epitopes. The eight groups are identified in the right panel of Figure 1(A). The top one shared motif refers to the motif that is shared by the largest number of epitopes in this group. 9/229 means 9 epitopes shared this motif in all 229 epitopes. (D) Root-mean-square deviations of 133 co-crystalized MHC-II-epitope complexes and grouped as three MHC-II types: DR (mean: 0.5581), DQ (mean: 0.8069), and DP (mean: 0.7985). (E) The structural alignment of 133 co-crystalized MHC-II epitopes on PDB as shown in sticks. (F) The structural alignment of five co-crystalized epitopes, 1bx2 (DR), 1klg (DR), 2nna (DQ), 4p5k (DP), and 5ujt (DQ) was used to number the positions. The eight most overlapped epitope positions, 1-6, 9, and 10, are shown as colorful sticks. (G) The left panel is the structural alignment of 1bx2 (magenta) and 3pl6 (red). The right panel is the structural alignment of 3pdo (magenta) and 3pgc (red), with a K(Lys) residue circled, indicating the start of the identical sequences. Dashed lines show the residue names of the sticks they pointed at.

We also performed the sequence alignments of the G-domains of 66 HLA class II alleles and found several conserved motifs shared among their binding grooves (Figure S2A). G-beta displayed more consecutive motifs compared to G-alpha. In G-alpha, alleles on the DRA1 locus have conserved motifs, like ‘FD’, shared with the DPA1 locus at the same position, while their corresponding positions are shifted three units to the right on DQA1 locus. In G-beta, the conserved motifs, like ‘RFDSDV’, of DRB1 are at the same position as DQB1, but they shift two units to the left in DPB1 (Figure S2A).

We then employed the MCCS scoring technique across the 133 crystalized MHC-II-epitope complexes (pMHC-IIs) to discern key residues in both the MHC-II binding groove and critical positions on the epitope sequence. Around 120 co-crystalized epitopes show high total energy contributions (< -0.7 kcal/mol) to the binding in positions 1-10 (Figure S2B, S2C). This is consistent with the results generated by Rosetta energy breakdown[4], which further confirmed the core binding regions on MHC-II epitope sequences (Figure S2D). Residues in this region can form steric interactions with MHC-II. In positions 1, 3, 5, 8, 9, and 10, residues interact with MHC-II via hydrogen bonds with a frequency more than 100 (Figure S2C). Only position 2 has more than 100 residues forming hydrophobic interactions with MHC-II (Figure S2C). Detailed residues’ interactions in DR, DP, and DQ groups can be found in Figure S3. As for the MHC-II G-domain, the scoring results revealed six conserved and six non-conserved residues with high total-energy contributions to the binding that shared among all 133 pMHC-IIs (Figure S2E, Figure S4). The conserved residues mainly interact with positions 1-10 via hydrogen bonds and steric interactions. A_PHE54 and B_TRP61/59 also engage in forming hydrophobic interactions (Figure S2F). Six non-conserved residues at corresponding positions within the G-domains of DR, DQ, and DP primarily influence binding at positions 1-10 (Figure S2F). These residues may be crucial for determining MHC-II epitope binding specificity. Detailed interactions of both conserved and non-conserved residues of G-domain are illustrated in Figure S5 and Table S1. The previously reported epitope’s anchor region that revealed by experimental structures, P1-P9 (PDB: 1J8H), here refers to the positions 2-10 in our case (PDB: 1bx2) can force peptide in the MHC-II binding groove by imposing hydrogen bonds from the conserved MHC-II residues[5, 6]. In many experimental structural analyses of pMHC-IIs, the interactions of anchor residues P1 (position 2), P4 (position 5), P6 (position 7), and P9 (position 10) received widespread attention[5, 6]. Position 2 favors hydrophobic interactions (Figure S2C). Positions 5 and 10 interact with negatively charged residues, Glu and Asp, which is consistent with the polar nature identified from the crystallography analysis of HLA-DQ2 (Table S1). Position 7 can form steric interactions and hydrogen bonds, for example, with positively charged residue Arg, Lys, and His (Table S1), which further convinced us that the computational results match those observed in the experimental structures[5, 6].

To summarize, while there is no distinct pattern in amino acid composition for antigenic epitopes bound by MHC-II at the sequence level, highly overlapping epitope backbones have been observed in the X-ray crystal structures of various pMHC-IIs. The residues at eight positions (1-6, 9, and 10) are also highly superimposed in their side chains. The G-domain sequences of DR-, DQ-, and DP-encoded MHC-II share multiple conserved motifs but show varying degrees of positional shifts across different types of MHC-II. Despite these shifts, the conserved motifs demonstrate a spatial overlap in aligned 3D structures. Among them, twelve (non-)conserved residues are identified to play critical roles in epitope binding within the MHC-II binding groove. Thus, we hypothesized that 3D structural features of epitopes are essential for identifying antigenic epitopes that gain the abilities for MHC-II binding and T-cell activation. Since different amino acids may form the same interactions if they have similar functional groups, it is possible to maintain or optimize the function of T cell activation by fixing the epitope in a similar 3D shape and designing the epitope’s sequence. However, one critical prerequisite is that the designed epitope sequences remain homologous to the antigen they derived from, ensuring the potential immune response can target the same site. This will be discussed later.

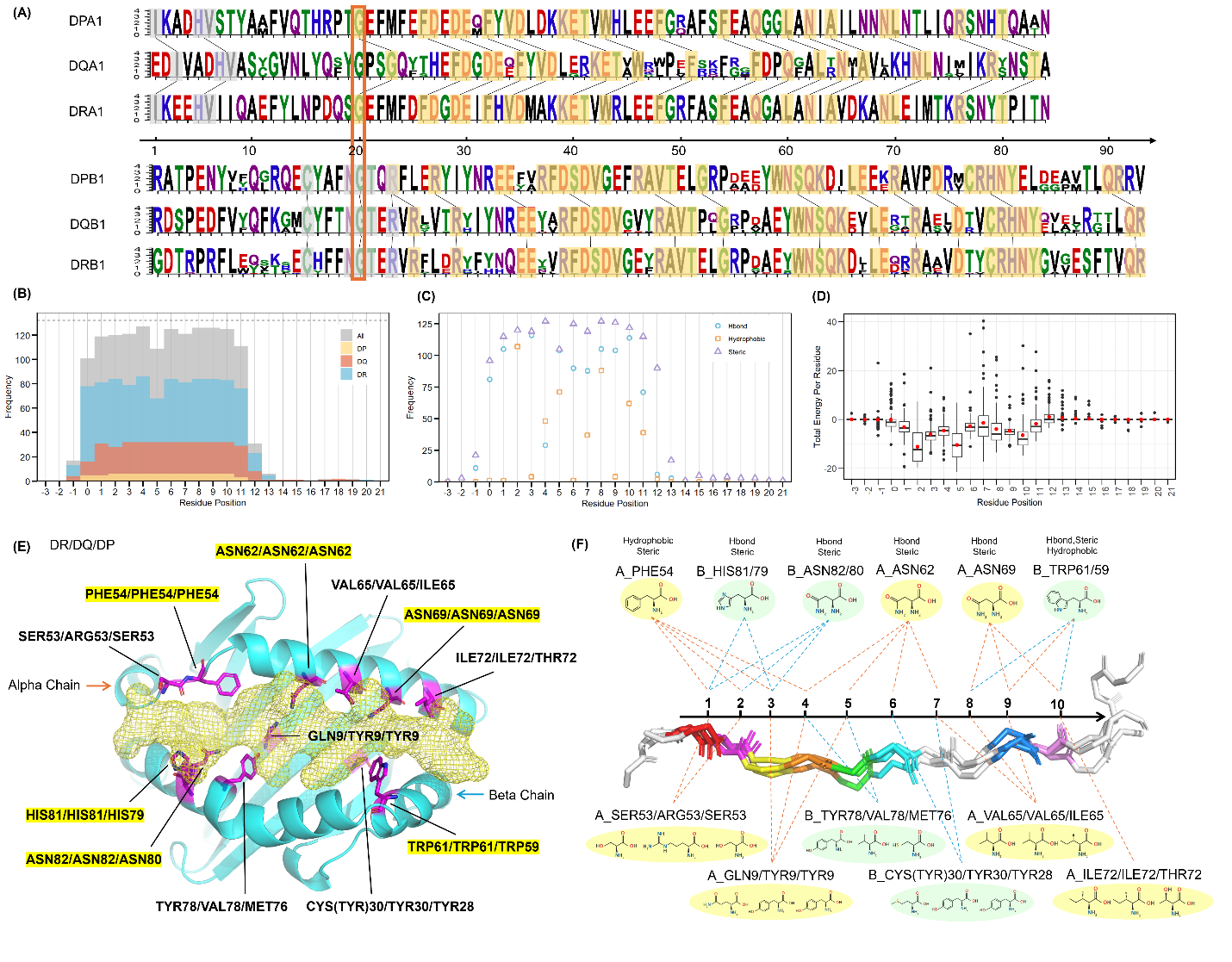

Figure S2 The detailed sequence and structure analysis of the key-binding residues and positions on MHC-II and epitopes, respectively. (A) Sequence logo plot of 66 HLA class II alleles with the conserved patterns among different types of MHC-II highlighted in yellow and gray. 66 alleles are divided into DPA1, DQA1, DRA1, DPB1, DQB1, and DRB1 groups. The x-axis shows the positions of the aligned sequences of the G-domain. Glycine with an orange square at position 20 indicates a different alignment pattern before and after this residue. (B) and (C) show MCCS scoring results of 133 co-crystalized epitopes using the position numbers in Figure 1(E). (B) The overlapped histograms of all 133, DR-bound, DQ-bound, and DP-bound epitopes indicate how many residues at each position have high-energy contributions (scores < -0.7 kcal/mol) to MHC-II binding. (C) The dots indicate how many residues significantly contribute (scores < -0.3 kcal/mol) to hydrogen bonds, hydrophobic interactions, or steric interactions at each position. (D) Each black dot represents the total energy for one residue at a specific position on the epitope sequences calculated by the Rosetta energy splitting algorithm using 103 MHC-II-epitope complexes with unmodified epitope sequences. The lower, the better. (E) The overview of the aligned residues on the MHC-II binding groove of DR/DQ/DP that highly contribute to the epitope binding calculated using MCCS. Conserved residues shared by all MHC-II are highlighted in yellow. The others are non-conserved residues. (F) A schematic diagram of MCCS results demonstrates which epitope positions interact with the conserved (above x axis) and non-conserved (below x axis) residues on the MHC-II binding groove. Dashed lines show the positions of the residues that interact with. The residues are shown as DR/DQ/DP.

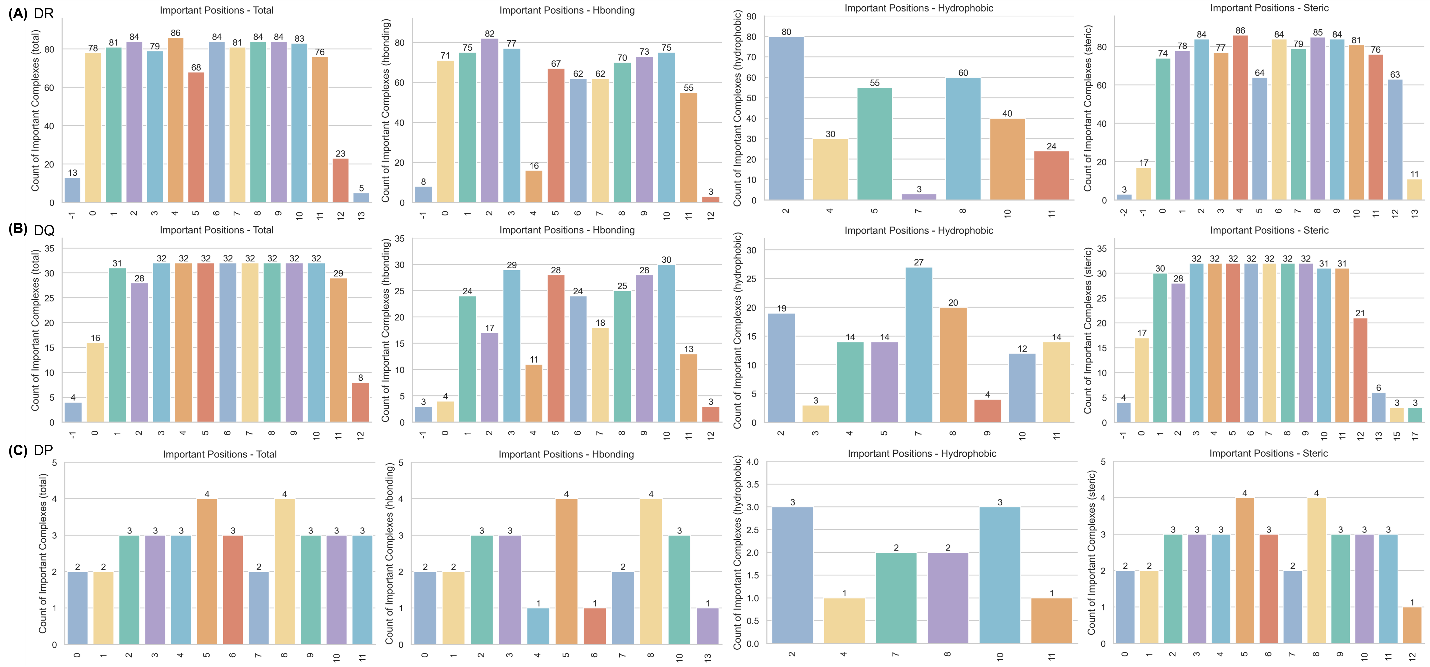

Figure S3 The MCCS scoring results of the 133 co-crystalized epitopes in (A) DR-epitope complexes (89), (B) DQ-epitope complexes (32), and (C) DP-epitope complexes (6), respectively, based on the previously defined position numbers in Figure 1(E). The results indicate how many residues in each position of epitopes highly contribute to the total binding to MHC-II (< -0.7 kcal/mol), to form hydrogen bonds (hbond) (< -0.3 kcal/mol), to form hydrophobic interactions (< -0.3 kcal/mol), and to form steric interactions (< -0.3 kcal/mol).

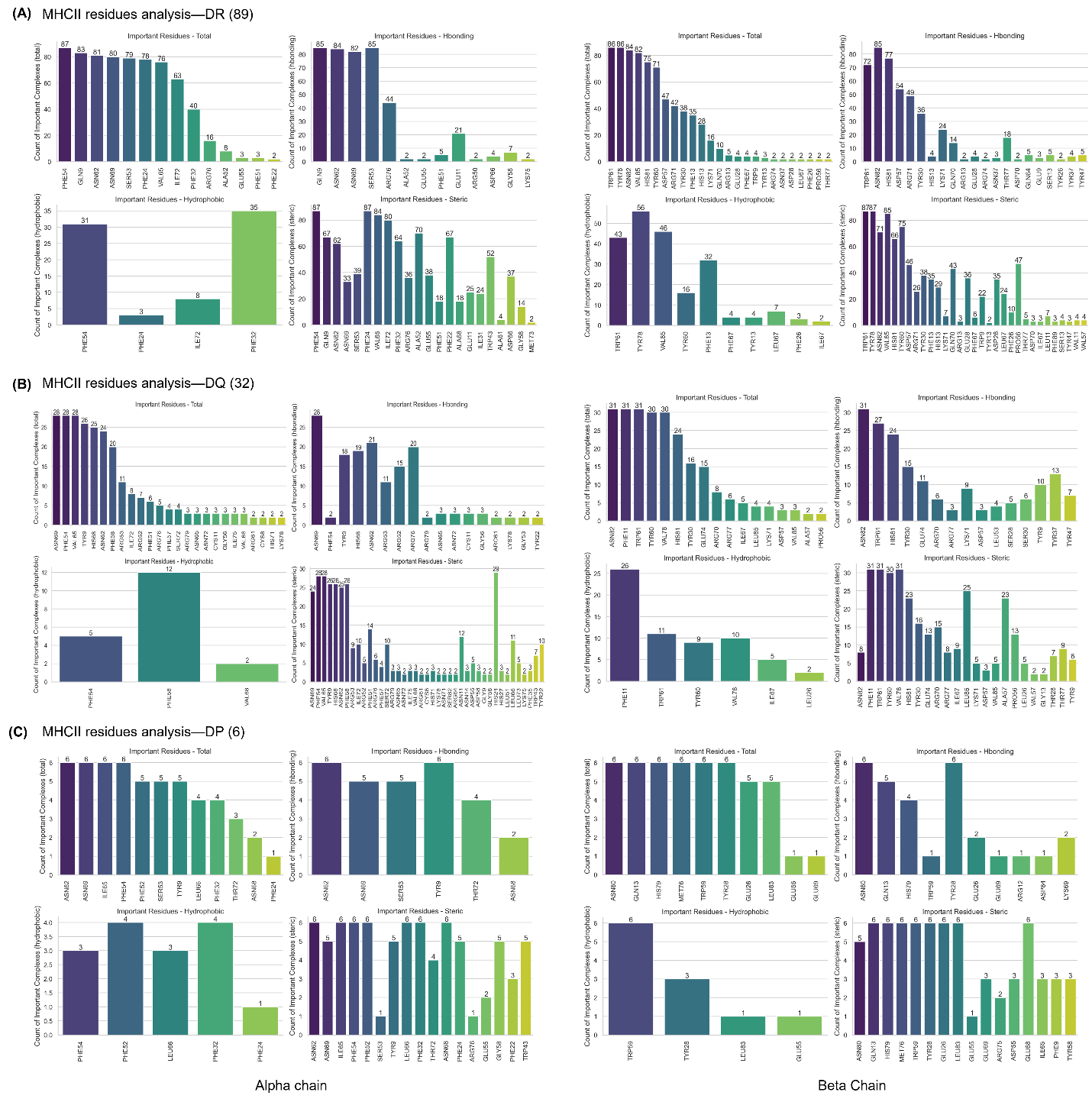

Figure S4 The MCCS scoring results of the 133 co-crystalized MHC-II G-domains in (A) DR-epitope complexes, (B) DQ-epitope complexes, and (C) DP-epitope complexes, respectively. The x-axis shows the residue's names. The y-axis is the count of complexes. The results indicate how many G-domain of the complexes have this residue that highly contributes to the total binding to its binding epitope (< -0.7 kcal/mol), to form hydrogen bonds (hbond) (< -0.3 kcal/mol), to form hydrophobic interactions (< -0.3 kcal/mol), and to form steric interactions (< -0.3 kcal/mol).

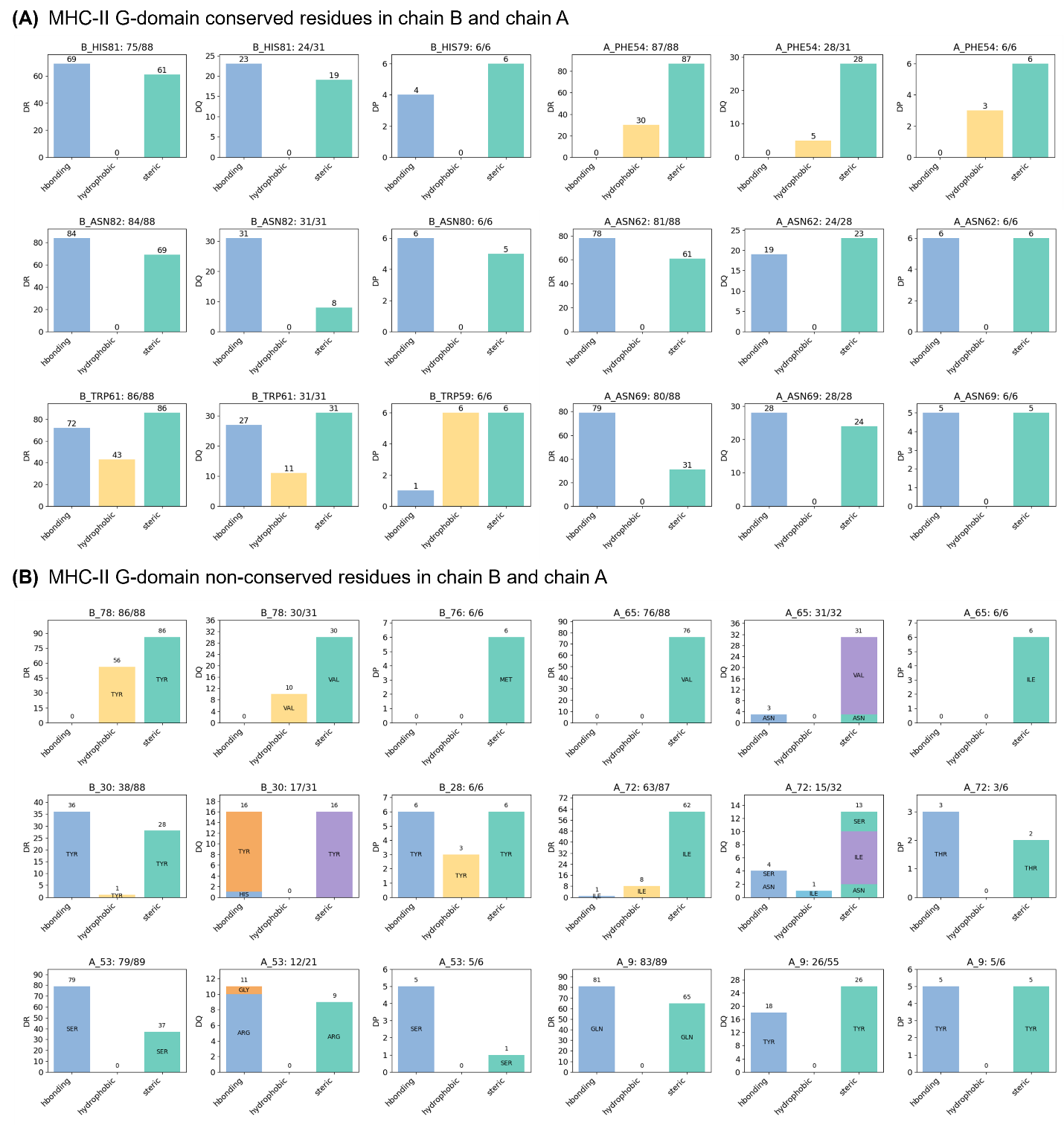

Figure S5 The detailed information on the interactions of the (A) conserved and (B) non-conserved residues on the G-domain. For example, ‘B_HIS81:75/88, B_HIS81: 24/31, B_HIS79: 6/6’ means the residue HIS81/79 is a conserved residue at the same positions in the DR, DQ, and DP-encoded MHC-II G-beta domains. For DR/DQ encoded MHC-II, HIS is numbered as 81, but for DP-encoded MHC-II, HIS is numbered as 79. ‘75/88’ means 75 DR-encoded MHC-II have this residue with total energy < -0.7 kcal/mol compared to 88 DR-encoded MHC-II with this residue in their sequences without missing (The total number of DR-encoded MHC-II is 89). The x-axis shows the types of interactions. The y-axis indicates the number of MHC-II that have this residue to form different types of interactions.

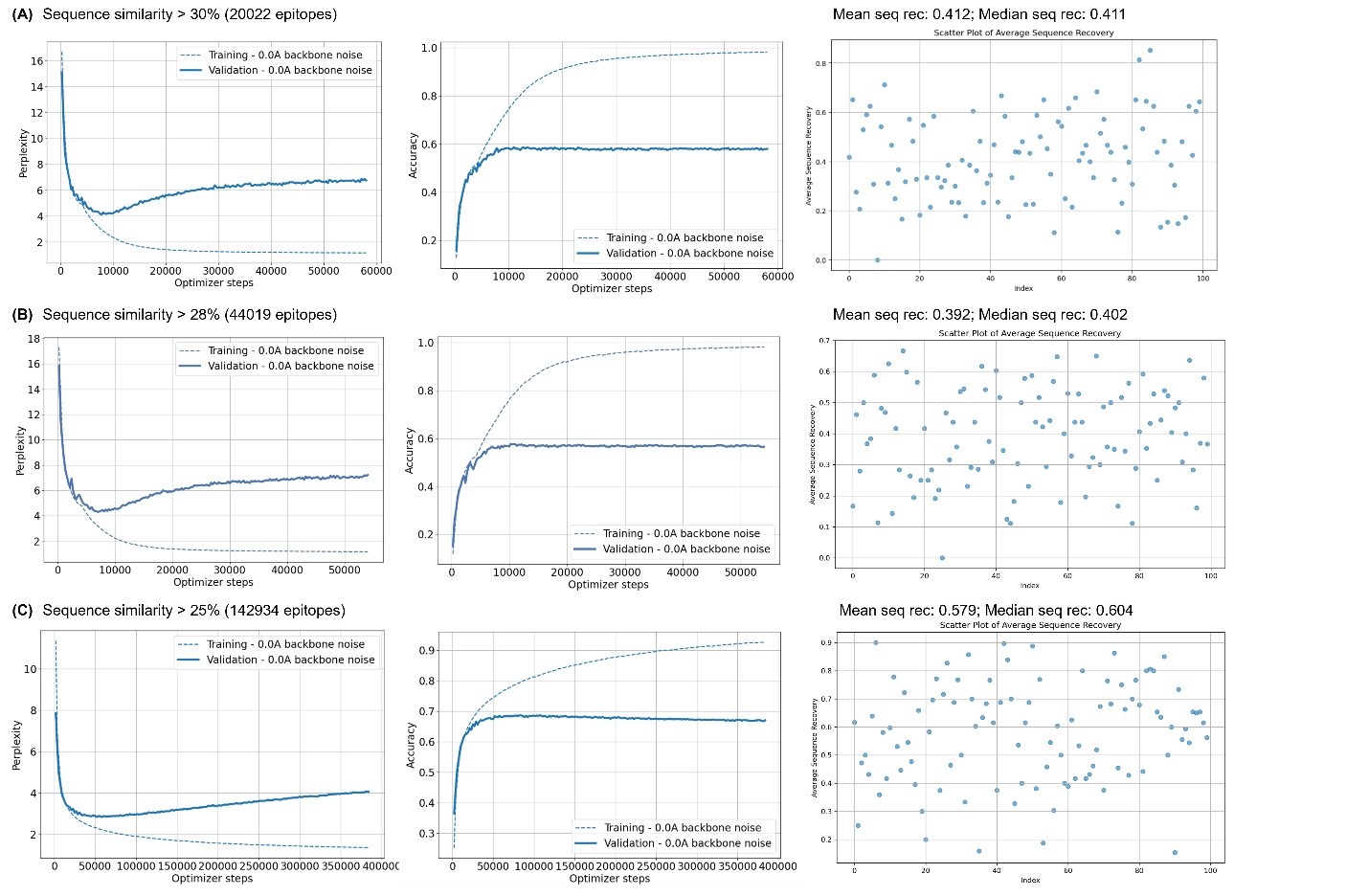

Figure S6 The simple ProteinMPNN model pre-training results. (A) The dataset which contains 20022 epitopes with sequence similarity larger than 30%. Training/Validation perplexity of the final epoch: 1.156/6.756; Training/Validation accuracy of the final epoch: 0.982/0.580. Mean sequence recovery: 0.412; Median sequence recovery: 0.411. (B) The dataset which contains 44019 epitopes with sequence similarity larger than 28%. Training/Validation perplexity of the final epoch: 1.156/7.218. Training/Validation accuracy of the final epoch: 0.982/0.565. Mean sequence recovery: 0.382; Median sequence recovery: 0.402. (C) The dataset which contains 142934 epitopes with sequence similarity larger than 25%. Training/Validation perplexity of the final epoch: 1.364/4.063; Training/Validation accuracy of the final epoch: 0.927/0/670. Mean sequence recovery: 0.579; Median sequence recovery: 0.604.

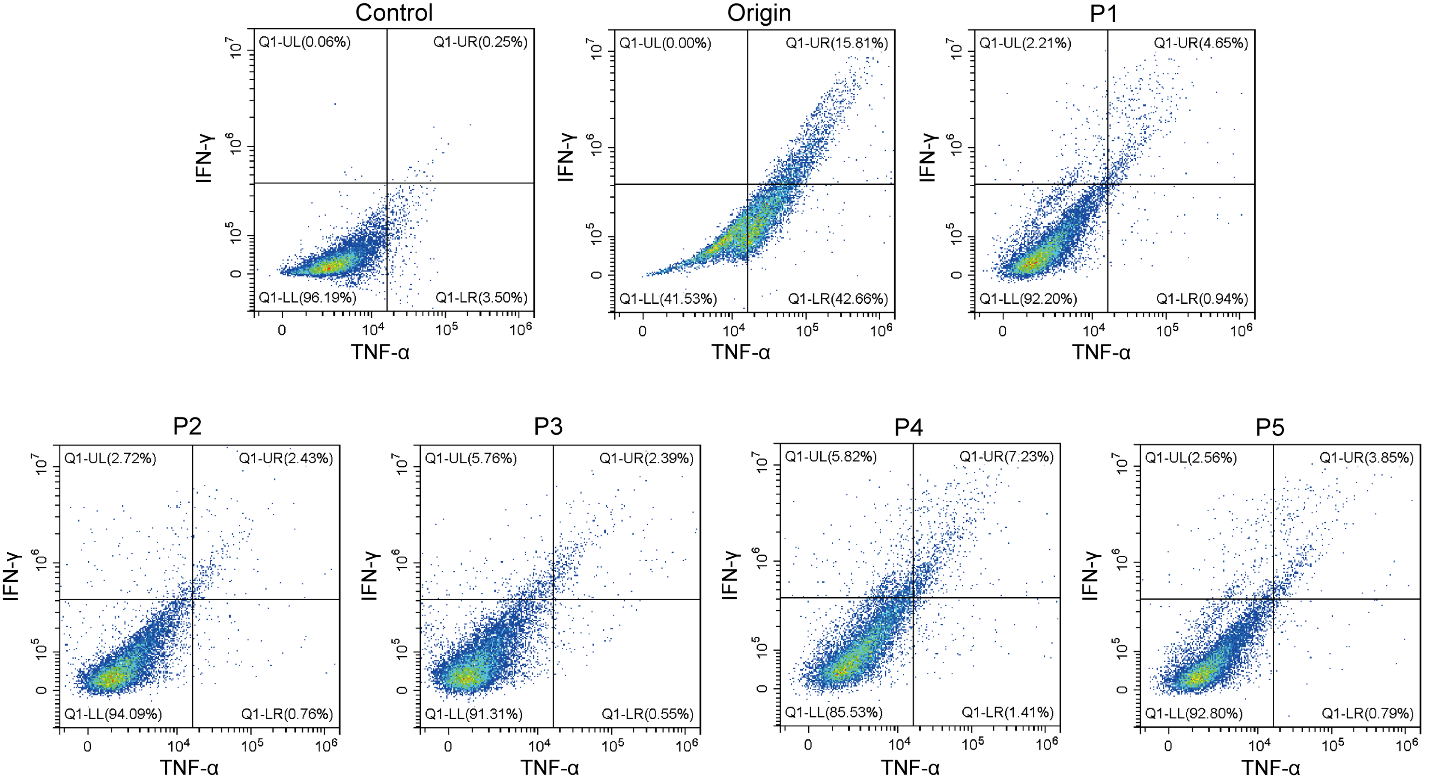

Figure S7 Secretion of IL-4 by CD4^+^ T cells in each group (IL-4 detected by APC).

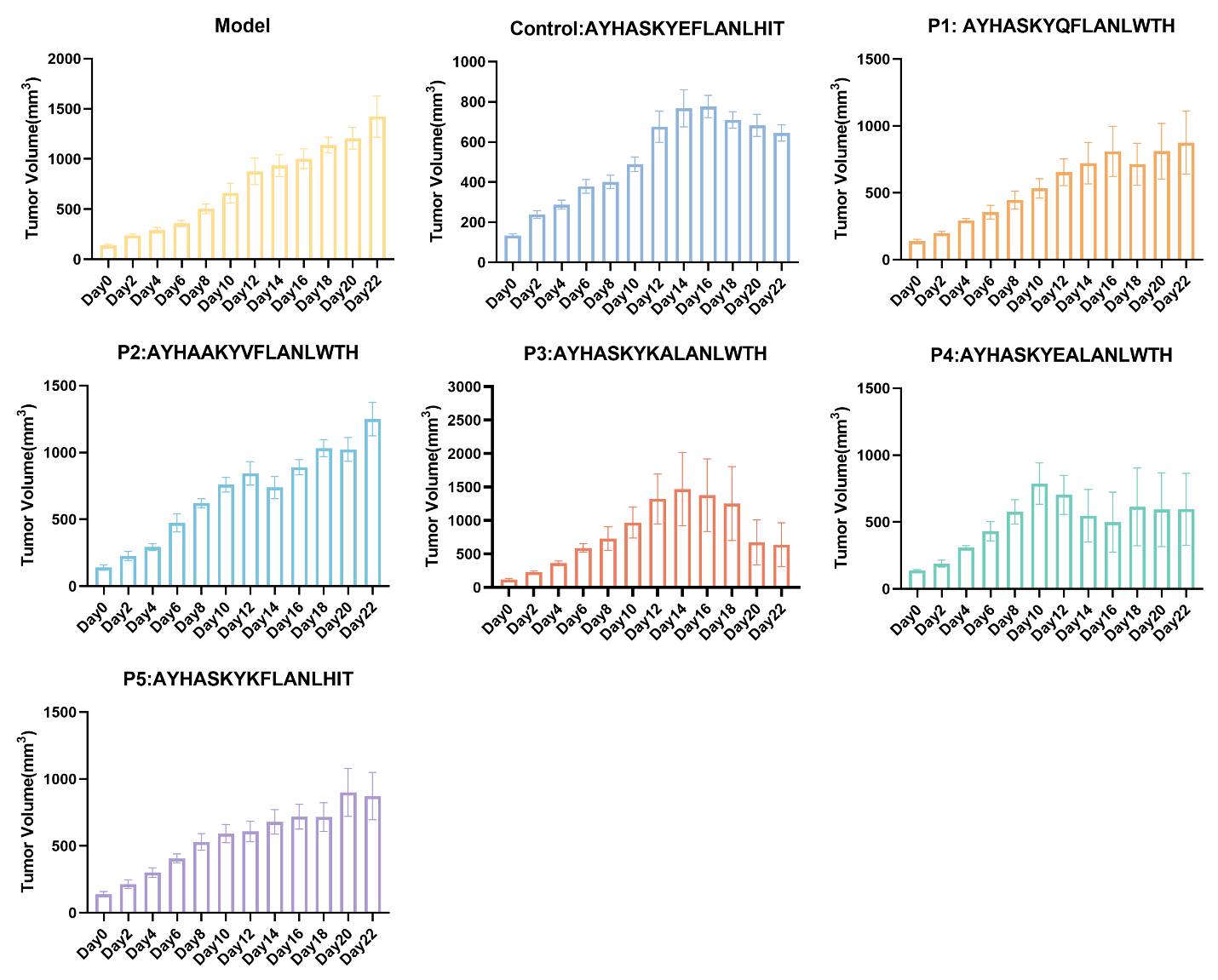

Figure S8 Tumor growth of mice in each group in 0-22 days. The error bar is shown as SEM.

The code for running MCCS scoring technique:

docker pull stcmz/mccs

docker run --rm -it -v "$(pwd):/data" stcmz/mccs

cd /data

chimera --nogui --script ~/incompleteSideChains.py 1aqd_MHC_II.pdb # Fix protein side chains.

vega 1aqd_MHC_II.pdb -o 1aqd_MHC_II.pdbqt -f VINA -c Gasteiger -p VINA -l GEN -r APOLAR -w # Convert MHC-II to PDBQT format.

vega 1aqd_epitope.pdb -o 1aqd_epitope.pdbqt -f VINA -c Gasteiger -p VINA -l GEN -r APOLAR -w # Convert epitope to PDBQT format.

pdbqtf $(grep --include=*.pdbqt -rl '?') -a # Fix problematic PDBQT files.

jdock -r 1aqd_MHC_II.pdbqt -l 1aqd_epitope.pdbqt -o output_MHCII -spa # Score the residues on the MHC-II binding groove.

jdock -r 1aqd_epitope.pdbqt -l 1aqd_MHC_II.pdbqt -o output_epitope -spa # Score the residues on the Epitope.
